# Prefrontal Cortex subregions provide distinct visual and behavioral feedback modulation to the Primary Visual Cortex

**DOI:** 10.1101/2024.08.06.606894

**Authors:** S. Ährlund-Richter, Y. Osako, K.R. Jenks, E. Odom, H. Huang, D.B. Arnold, M. Sur

## Abstract

The mammalian Prefrontal Cortex (PFC) has been suggested to modulate sensory information processing across multiple cortical regions via long-range axonal projections. These axonal projections arise from PFC subregions with unique brain-wide connectivity and functional repertoires, which may provide the architecture for modular feedback intended to shape sensory processing. Here, we used axonal tracing, axonal and somatic 2-photon calcium imaging, and chemogenetic manipulations in mice to delineate how projections from the Anterior Cingulate Cortex (ACA) and ventrolateral Orbitofrontal Cortex (ORB) of the PFC modulate sensory processing in the primary Visual Cortex (VISp) across behavioral states. Structurally, we found that ACA and ORB had distinct patterning of projections across cortical regions and layers, monosynaptically targeting both excitatory and inhibitory neuronal cell types in these areas. ACA axons in VISp had a stronger representation of visual stimulus information than ORB axons, but both projections showed non-visual, behavior-dependent activity. ACA input to VISp enhanced the encoding of visual stimuli by VISp neurons, and modulation of visual responses scaled with arousal. On the other hand, ORB input shaped movement and arousal related modulation of VISp visual responses, but surprisingly reduced the encoding of high-contrast visual stimuli. Thus, ACA and ORB feedback have separable projection patterns and encode distinct visual and behavioral information, putatively providing the substrate for their unique effects on visual representations and behavioral modulation in VISp. Our results offer a refined model of cortical hierarchy and its impact on sensory information processing, whereby specific PFC projections contribute uniquely to VISp activity during discrete behavioral states.

## Introduction

A long-long standing hypothesis of mammalian brain organization suggests that brain regions are hierarchically organized, and that the relative hierarchical order of two regions can be identified by the laminar connectivity between them^1,2^. This bidirectional connectivity is proposed to allow higher order regions to influence, or modulate, the activity of lower cortical regions to optimize information processing, particularly sensory information^3,4^. Large-scale connectivity mapping of the mouse brain has revealed that the Prefrontal Cortex (PFC) and its subregions constitute the top of the cortical hierarchy, and that the majority of PFC efferent axons are classified as feedback projections^5^. Discrete subregions of the PFC target both shared and unique cortical regions^6,7^, and combined, PFC neurons reach all of the cortex via a sophisticated pattern of axonal innervation^8,9^. This anatomical organization ideally positions the PFC to guide or modulate cortex-wide activity and thereby shape information processing and ultimately behavior. Indeed, perturbation of PFC activity has been observed to not only disrupt cortex-wide activity dynamics, but also behavioral performance^10,11^. However, it remains unclear whether this sophisticated anatomical heterogeneity of projection populations within the PFC is matched by equivalently heterogenous functional feedback, or if the PFC provides more general brain-state dependent feedback cortex-wide.

A key feedback circuit, present across species and well-studied in rodent models, is the cortical projections from the PFC to the Primary Visual Cortex (VISp)^12,13^. In the mouse, the VISp is targeted by two distinct PFC subregions, the Anterior Cingulate Cortex (ACA) and the ventrolateral Orbitofrontal Cortex (ORB)^6,14–16^. Previous work has suggested a functional dichotomy of ACA and ORB feedback to the VISp during visually-guided behavioral tasks. The ACA has been suggested to enhance the encoding of visually relevant stimuli^17^, i.e., to facilitate selective visual attention, while the ORB has been suggested to filter out non-relevant visual information to enable associative learning^18^. Two, opposing local inhibitory microcircuit circuit motifs have also been described, where ACA axons recruit VIP positive interneurons to enhance the response amplitude of VISp neurons^17^, while ORB axons instead recruit SST positive interneurons to reduce the response amplitude of VISp neurons^18^. These studies have offered a circuit-based mechanism as to how feedback modulation could alter the task-dependent representation of visual stimuli depending on the level of behavioral relevance (i.e., rewarded/ unrewarded). However, the activation of these feedback projections by optogenetic constructs provides an artificial pattern of activity, and their physiological relevance in relation to more natural behavioral states still remain unclear.

Recent studies on the complexity of PFC axonal innervation of the cortex has revealed that both the ACA and the ORB have projection neurons that can selectively target visual regions, and also that these neurons have more extensive collateral projections reaching other cortical modalities, e.g., the primary Motor Cortex (MOp), in addition to visual regions^8,9^. This discovery has created a vacuum in our understanding of PFC feedback that is shared across regions, and modulation that is specific towards a discrete sensory modality. As a consequence, microcircuitry targeted by and functional content of these collaterals in distinct cortical regions remains unexplored.

Here, we sought to clarify the unique information that ACA and ORB axons convey to the VISp during natural behaviors, and what effects these inputs have on the encoding of visual stimuli by VISp neurons. By imaging the activity of ACA and ORB axons in both the VISp and the MOp, we observed a stronger representation of visual information in ACA axons in comparison to ORB axons, and a stronger representation of movement velocity in the MOp in comparison to the VISp. ACA axon activity was modulated in graded manner by arousal. Using chronic imaging of VISp neurons with and without chemogenetic suppression of ACA or ORB feedback, we found that the ACA and ORB play complementary roles in the modulation of VISp. ACA inputs enhance visual encoding in VISp during arousal states. ORB inputs link high arousal and movement modulation to VISp activity even at the expense of visual encoding. Our data support a model of PFC feedback that is specialized at both the level of PFC subregions and their targets, enabling each region to selectively shape target-specific cortical activity rather than modulating it globally.

## Results

### Cortical targets of ACA and ORB projection neurons.

As a first step in characterizing ACA and ORB feedback projections, we mapped the axonal density and the cellular identity of post synaptic neurons in downstream cortical regions, with a particular focus on the VISp and MOp. To verify stereotaxic coordinates for targeting VISp-projecting neurons, we injected retroAAV-hSyn-GFP into VISp and identified dense labeling in ACA and ORB (Supplementary Figure 1a–d), consistent with prior work extensively characterizing these projections^19^ (**Supplementary Figure 1A-D**). To carefully map the density of axons in cortical regions targeted by projection neurons within the ACA and the ORB, AAV-CAG-tdTomato and AAV-CAG-GFP were injected unilaterally into the ACA and the ORB, respectively (**Figure 1A and Supplementary Figure 1E**). To generate a cortex-wide map of ACA and ORB axonal projections, selected brains were cut coronally (50μm), and every other section was mapped to the Allen Reference Atlas (ARA)^20^ using a modified version of the WholeBrainSuite in R^20^, resulting in a density estimate (μm^2^_axon_/μm^2^_total area_) of axonal projections for each cortical region and layer (as defined by the Allen Brain Atlas) (**Figure 1B**).

**Figure 1.**
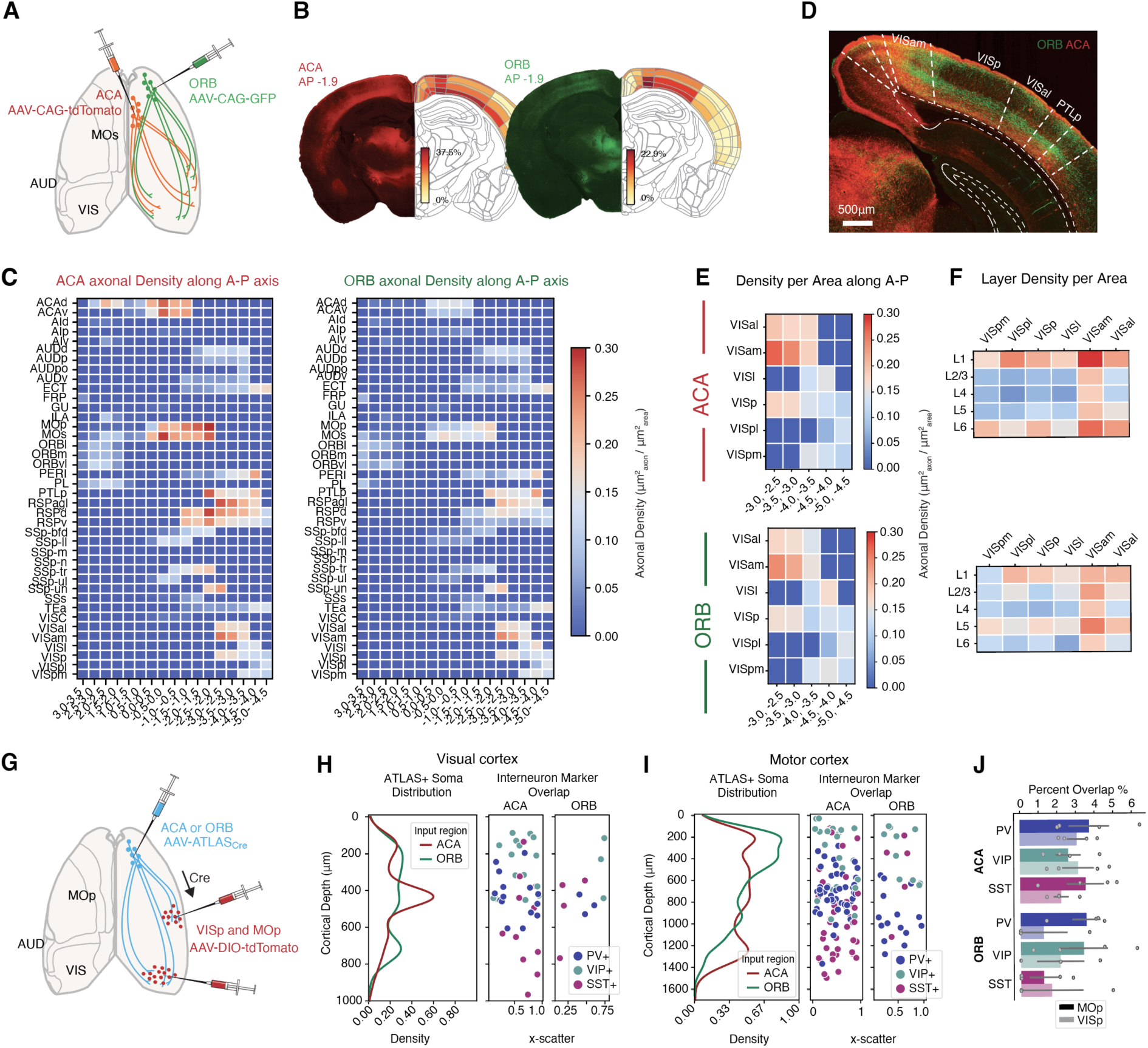
Axonal projections from the ACA and the ORB innervate discrete layers of the VISp and adjacent regions, and target diverse neuron classes. **(A)** Experimental strategy. AAV-CAG-tdTomato and AAV-CAG-GFP was unilaterally injected into the ACA and ORB, respectively. **(B)** Example image of a brain section (with the green and red channel split) mapped to the Allen Brain Atlas and the axonal density of each cortical region and layer measured for that section. **(C)** Average axonal density per cortical region and anterior-posterior (A-P) coordinate bin over all mice (n=4, 67±20 brain sections per animal) from the ACA (left) and ORB (right). **(D)** Example image of primary visual cortex (VISp) and surrounding visual regions with ORB axonal innervation (green) and ACA axonal innervation (red) targeting superficial and deep layers. (Split channels of image in Supplementary Figure 1). **(E)** Average axonal density of ACA (top) and ORB (bottom) projections in the visual regions of the mouse cortex along the A-P axis (n=4 mice). **(F)** Axonal density per cortical layer for all visual regions of the mouse cortex (n=4 mice). **(G)** Experimental strategy. AAV-ATLAS_Cre_ was unilaterally injected into the ACA or the ORB, respectively, followed by an injection of AAV-DIO-tdTomato into the VISp and MOp. **(H)** Left, Kernel Density Estimate (KDE) density distribution of tdTomato+ neurons monosynaptically targeted by either the ACA (red) or ORB (green) along the depth of the visual cortex. Right, tdTomato+ neurons overlapping with immunohistological labeling of PV (blue), SST (purple) or VIP (green) along the depth of the visual cortex, all analyzed sections superimposed along AP axis. ACA: n= 4 mice, MOp n=32 sections, VISp n=31 sections. ORB: n= 3 mice, MOp:30 sections, VISp: 15 sections). **(I)** Left, KDE density distribution of tdTomato+ neurons monosynaptically targeted by either the ACA (red) or ORB (green) along the depth of the motor cortex. Right, tdTomato+ neurons overlapping with immunohistological labeling of PV (blue), SST (purple) or VIP (green) along the depth of the motor cortex, all analyzed sections superimposed along AP axis. ACA: n= 4 mice, MOp n=32 sections, VISp n=31 sections. ORB: n= 3 mice, MOp:30 sections, VISp: 15 sections). **(J)** Percentage overlap of tdTomato+ neurons with immunohistological labeling of PV (blue), SST (purple) or VIP (green) in MOp (dark) and VISp (light). (ACA: n= 4 mice, MOp n=32 sections, VISp n=31 sections. ORB: n= 3 mice, MOp:30 sections, VISp: 15 sections). Error bars: SEM, dots represent individual animals.

ACA and ORB projection neurons broadly targeted largely overlapping cortical regions, such as visual, motor, somatosensory and auditory cortical regions, as described previously^7^, although the density of axonal projections within each target differed between the two PFC source regions (**Figure 1C**, and **Supplementary Figure 1F-I**). The axonal mapping revealed dense axonal projections from both the ACA and the ORB innervating Retrosplenial (RSP) and Visual areas, but while the ACA targeted both the dorsal and ventral RSP, the ORB more restrictively targeted the dorsal RSP. The two PFC regions innervated all of the major visual cortical regions, with the highest axonal density present in the anteromedial visual cortex (VISam), where the axonal projections from both regions innervated all layers of VISam (**Figure 1D-F**). In the rest of the visual regions (VISal, VISl, VISp, VISpl, VISpm) ACA and ORB axonal projections displayed distinct innervation patterns, where ACA axons discretely targeted layers 1 and 6, while ORB axonal projections were specific to layers 1 and 5 (**Figure 1F**, and **Supplementary Figure 1J**). Notably, while the differences between ACA and ORB axonal innervation of the layers of VISp has been documented before^16^, our cortex-wide axonal mapping showed that the distinct patterns of innervation from the ACA and ORB were present across all visual cortex regions, while they were not particularly distinct in other sensory or motor regions (**Supplementary Figure 1F and H**).

To further investigate the cellular targets of ACA and ORB projections in VISp and MOp, we utilized the Anterograde Transsynaptic Label based on Antibody-like Sensor (ATLAS) viral tracer injected into ACA or ORB to deliver Cre to post synaptic neurons^21^. Subsequently, a viral injection of AAV-DIO-tdTomato in the VISp and MOp allowed for the visualization of neurons monosynaptically targeted by either ACA or ORB projections (ACA: n=4 mice, ORB: n=3 mice, **Figure 1G**, and **Supplementary Figure 1K**). This approach was particularly valuable in the context of long-range PFC inputs, which target dendrites in superficial layers (notably Layer 1), while the somata of their postsynaptic partners may reside in deeper layers such as Layer 5^22^. Thus, axonal projection density alone does not necessarily predict the laminar location of postsynaptic cell bodies. Immunohistochemical labeling of three major interneuron classes—Somatostatin (SST), Parvalbumin (PV), and Vasoactive Intestinal Peptide (VIP)—revealed that both putative excitatory (triple-negative) and inhibitory neurons are monosynaptically targeted by ACA and ORB axons in both VISp and MOp across all layers (**Figure 1H-J, Supplementary Figure 1L**).

Overall, we found that long-range projections of ACA and ORB target discrete layers in visual cortex and have access to both excitatory and inhibitory local microcircuitry. This anatomical difference of connectivity between the two PFC regions thus suggests that differences may exist in their influences on VISp activity.

### ACA axons have stronger visually driven activity that scales with contrast, compared to ORB axons

To assess the modulatory activity transmitted from the ACA and the ORB to the VISp, we performed 2-photon imaging of ACA or ORB axonal calcium activity ranging from 10 to 150μm in depth. As PFC projection neurons are known to have extensive collaterals to multiple cortical regions^9^, ACA and ORB axonal calcium activity in the MOp was also recorded in a separate cohort of mice, allowing for the comparison of modulatory activity putatively transmitted on a cortex-wide scale, to modulatory activity unique to the VISp or MOp. ACA or ORB axons were transduced with AAV-axon-GCaMP6s to allow for the calcium indicator to be expressed in the axons of ACA or ORB neurons. A cranial window was placed above the VISp or MOp and axonal activity at single-bouton resolution was recorded in layer 1 and superficial layer 2 (**Figure 2A-C**). Once habituated to the 2-photon microscope set-up and head-fixation, awake behaving mice were free to run on a running-wheel while visual stimuli were presented to the contralateral eye (in relation to the cranial window, **Figure 2D**). All mice experienced two different types of visual stimuli sessions each day, one consisting of repeated presentation of drifting gratings of 8 different directions at 3 different contrasts, and another with repeated presentation of five natural movie scenes at 3 different contrasts (see Methods for details). For some sessions, un-cued air puffs were delivered to the ipsilateral eye and facial area on alternate blocks of trials to elicit changes in arousal (see Methods for details). Active axonal segments (regions of interests (ROIs)) were identified using Suite2p^23^. To avoid artificial correlations of boutons from the same axonal branch, identified ROIs were clustered based on activity correlation measurements and activity traces of boutons from the same putative axons were averaged together to represent one axonal branch^24^ (see Methods). An axon was considered visually responsive if the mean ΔF/F activity while a stimulus was on was significantly different from the mean ΔF/F activity in the one second preceding visual stimuli onset for at least one type (one direction/movie) of stimuli (Student’s paired t-test, p-value <0.00625 for gratings, and p-value <0.001 for movies, **Figure 2E**). ACA axons in the VISp had the largest proportion of visually responsive axons (gratings: 0.17±0.03, 122±14 responsive axons, movies: 0.19±0.02, 125±9 responsive axons, mean±SEM,) followed by ACA axons in the MOp (gratings: 0.13±0.03, 182±57 responsive axons, movies: 0.13±0.01, 119±56 responsive axons, n=4 mice). The proportion of visually responsive ORB axons in the VISp (gratings: 0.06±0.01, 31±25 responsive axons, movies: 0.13±0.02, 42±29 responsive axons) and MOp (gratings: 0.07±0.01, 84±17 responsive axons, n=4 mice, movies: 0.11±0.01, 123±41 responsive axons,) was significantly lower than for ACA axons in the VISp (Gratings: ACA-VIS vs ORB-VIS: p-value= 0.0345, ACA-VIS vs ORB-MOp: p-value= 0.0231. Movies: ACA-VIS vs ORB-MOp: p-value= 0.0432, one-way ANOVA with post-hoc Tukey test for multiple comparisons of means.) (**Figure 2F-G)**. The majority of visually responsive axons from both the ACA and ORB exhibited increased activity in response to visual stimuli (**Supplementary Figure 3A–B**) and were selective for a specific direction of drifting gratings or a particular movie identity (**Figure 2F-G**).

**Figure 2.**
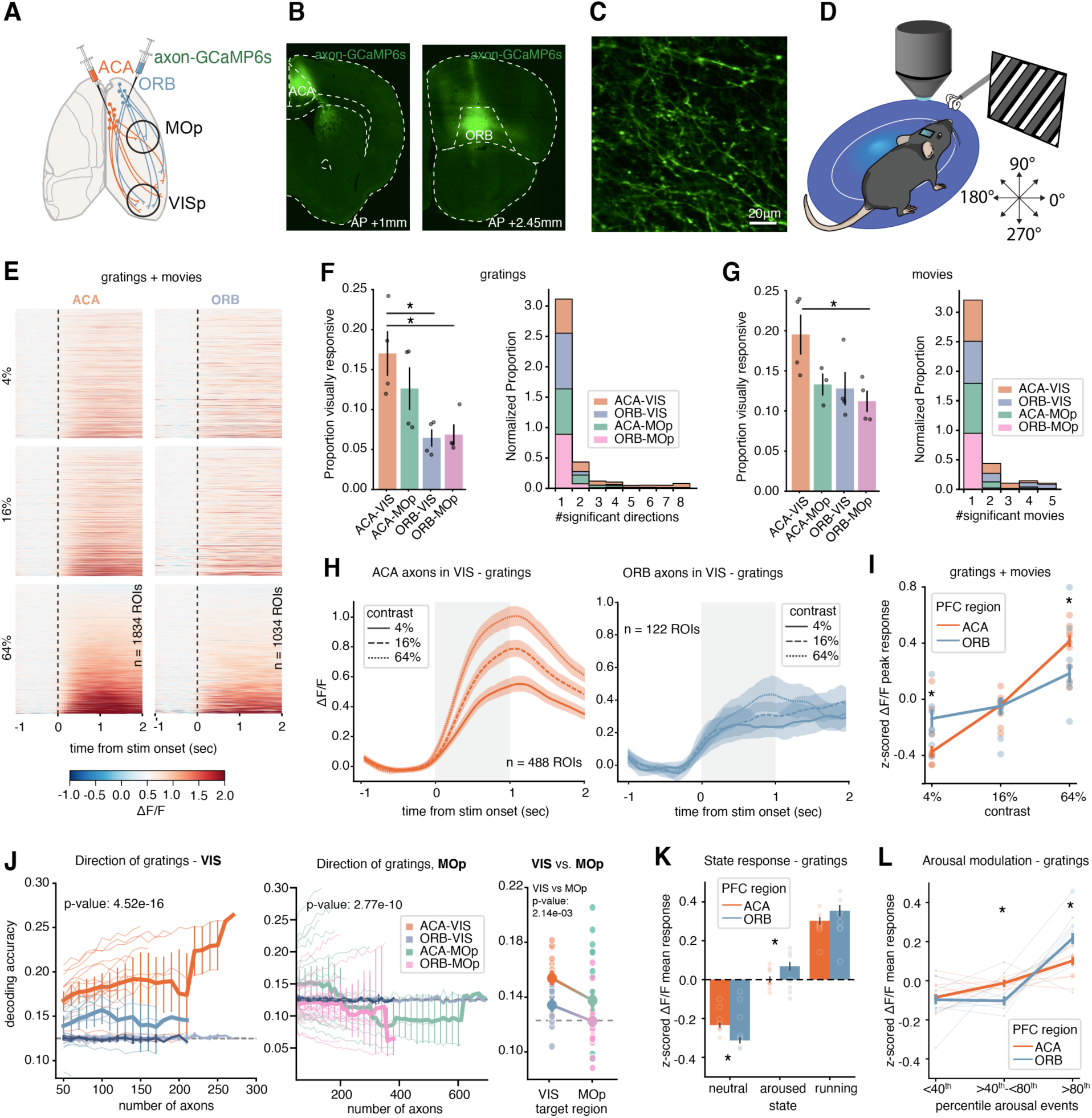
ACA and ORB axonal activity in VIS and MOp convey visually driven information that is differently modulated by behavioral state. **(A)** Experimental strategy. Axon-targeting GCaMP6s (AAV-hSyn-axon-GCaMP6s) was injected unilaterally in either the ACA or the ORB and a cranial window was implanted above either the VIS or the MOp. **(B)** Representative images of viral injection sites in the ACA (left) and the ORB (right). Replicated in n=4 ACA-VIS, n=4 ORB-VIS, n=4 ACA-MOp, and n=4 ORB-MOp mice. **(C)** Example field of view from 2-photon imaging of ACA axons in VIS. **(D)** Schematic of behavioral setup. **(E)** Heatmap of visually responsive axons in the MOp and VIS from the ORB (left) or ACA (right). Average responses across ten trials for each contrast and direction/movie, responses to all directions/movies are included. Axons are sorted by peak response amplitude to 64% contrast. ACA – VIS and MOp: 1834 axons from 8 mice, ORB– VIS and MOp: 1034 axons from 8 mice. **(F)** Proportion of visually responsive axons to gratings (left) from the ACA or the ORB in the VIS or the MOp. Bars display mean of proportions, error bars: SEM, dots represent individual animals. ACA-VIS vs ORB-VIS: p-value: 0.0345, ACA-VIS vs ORB-MOp: p-value: 0.0231, one-way ANOVA with post-hoc Tukey test for multiple comparisons of means. Selectivity of visually responsive axons (right), from the ACA or the ORB in the VIS or the MOp, shown as the normalized distribution of the number of directions to which each axon responded significantly. **(G)** Proportion of visually responsive axons to movies (left) from the ACA or the ORB in the VIS or the MOp. Bars display mean of proportions, error bars: SEM, dots represent individual animals. ACA-VIS vs ORB-MOp: p-value= 0.0432, one-way ANOVA with post-hoc Tukey test for multiple comparisons of means. Selectivity of visually responsive axons (right), from the ACA or the ORB in the VIS or the MOp, shown as the normalized distribution of the number of movies to which each axon responded significantly. **(H)** Averaged population responses of all visually responsive axons with an increase in ΔF/F in response to gratings, plotted per contrast (left; ACA-VIS axons, right; ORB-VIS axons). ACA-VIS 488 axons from 4 mice, ORB-VIS 122 axons from 4 mice. Solid line: mean value, shaded area: 95% confidence interval. **(I)** Standardized ΔF/F peak responses during stimuli on-time plotted per contrast. Visually responsive axons in VIS and MOp, with a significant increase in ΔF/F in response to stimuli are included. ΔF/F peak responses are z-scored across all trials and contrasts for each axon, such that negative values reflect responses lower than the axon’s average, while positive values indicate above-average amplitudes. ACA – VIS and MOp: 1854 axons, ORB– VIS and MOp: 1053 axons. Solid line: mean standardized ΔF/F peak responses, error bars: 95% confidence interval, dots: individual animals. ACA vs ORB: 4% p-value: 0.001, 64% p-value: 0.001, one-way ANOVA with post-hoc Tukey test for multiple comparisons of means. **(J)** Decoding accuracy of SVM decoder of drifting grating directions trained and tested on ACA (orange/green) or ORB (blue/pink) axonal responses (mean ΔF/F during stimuli on-time) in the VIS (left) or MOp (middle), per imaging session and field of view. Decoding accuracy of shuffled ACA axonal responses (light gray) or shuffled ORB axonal responses (dark gray) plotted as comparison. Solid line: mean accuracy across sessions, error bars: 95% confidence interval (omitted when only one session contributed to a bin), lines: individual imaging sessions, dashed line: chance level. p-value from 2-way ANCOVA to examine the effects of subset (number of axons) and PFC subregion (ACA vs. ORB) on the accuracy of decoding. The model included interaction terms to explore if the effect of subset on accuracy varies by group and data/shuffled data. p-value from the variability in accuracy explained by group (ACA vs. ORB). VIS-ACA vs. ORB: p-value: 4.52e-16, MOp-ACA vs. ORB: p-value: 2.77e-10. Average decoding accuracy across sessions in VIS and MOp of both ACA and ORB axons (right). Each dot represents one session, with color coding distinguishing between the four groups: ACA_MOp, ACA_VIS, ORB_MOp, and ORB_VIS. Independent samples t-tests were used to compare accuracy between MOp and VIS, p-value: 2.14e-3. VIS-ACA: 18 fields of view across 4 mice, ORB: 12 fields of view across 4 mice. MOp-ACA: 21 fields of view across 4 mice, ORB: 26 fields of view across 4 mice. **(K)** Standardized ΔF/F mean responses during stimuli on-time of visually responsive axons in VIS and MOp, with an increase in ΔF/F in response to grating stimuli, per behavioral state. Behavioral state was classified as the >80^th^ percentile trials per session (running speed: movement, pupil size: arousal, <80^th^ percentile events: neutral). Axonal responses to a direction were only included in the analysis if there were trials representing all three states. ΔF/F mean responses are z-scored across each axon. Bars: mean standardized ΔF/F mean responses across axons, error bars: 95% confidence interval, dots: individual animals. ACA vs ORB: neutral p-value: 0.001, aroused p-value: 0.001, one-way ANOVA with post-hoc Tukey test for multiple comparisons of means. **(L)** Standardized ΔF/F mean responses during stimuli on-time of visually responsive axons in VIS and MOp, with an increase in ΔF/F in response to grating stimuli, per arousal state. Arousal states was classified as: a pupil size z-score within each trial <40^th^ percentile pupil size (neutral), >40^th^ - <80^th^ percentile pupil size, and >80^th^ percentile pupil size, per session. Axonal responses to a direction were only included in the analysis if a drifting grating direction was presented at least once during all each arousal state. ΔF/F mean responses are z-scored across each axon. Solid line: mean standardized ΔF/F mean responses across axons, error bars: 95% confidence interval, thin lines and dots: individual animals. ACA vs ORB: >40^th^ - <80^th^ percentile p-value: 0.001, >80^th^ percentile p-value: 0.0061, one-way ANOVA with post-hoc Tukey test for multiple comparisons of means.

The visual responses of excitatory neurons in visual areas scaled with the contrast of the stimulus^25^. Notably, the mean and peak amplitude of response of the visually responsive ACA axons in both the VISp and MOp also scaled with the contrast of the visual stimuli presented, while contrast scaling was almost absent in ORB axonal activity (**Figure 2H**, and **Supplementary Figure 2**). ACA axonal activity in response to both drifting gratings and natural movies showed a greater difference in response to the highest and lowest contrast, in comparison to ORB axons (z-scored peak ΔF/F value for visually responsive axons and the stimulus/stimuli they were significantly responsive towards, contrast: 4%, p-value <0.001, contrast: 16%, p-value >0.05, contrast: 64%, p-value <0.001, one-way ANOVA with post-hoc Tukey test for multiple comparisons of means) (**Figure 2I**).

While a higher proportion of ACA axons are reliably driven by visual stimuli, it is possible that the population activities of ACA and ORB axons contain similar predictive information of the stimulus identity. To test this, we trained a linear support vector machine (SVM) to decode the identity of the visual stimulus from the mean ΔF/F responses of axonal activity in VISp in response to discrete stimuli. The decoder trained on ACA axon activity outperformed the decoder trained on ORB axonal responses (2-way ANCOVA, C(group, ACA vs. ORB), Direction: p-value = 4.52e-16, **Figure 2J**, left). While decoding accuracy based on mean responses did not differ between ACA and ORB axons for movie identity, temporal decoding accuracy was enhanced in ACA axons for both natural movies and gratings in VISp (**Supplementary Figure 3C-D**). Direct comparison of the predictive information regarding stimulus identity in PFC axon activity projecting to the VISp versus MOp revealed that both ACA and ORB transmit stronger visual information to VISp (**Figure 2J)**. Specifically, decoding accuracy of PFC axons was higher in VISp compared to MOp (Independent samples t-tests, VISp vs. MOp: p-value= 2.13e-3). These results suggest that visual information is encoded more robustly in ACA axons than in ORB axons within their respective downstream targets, with both PFC regions transmitting more visual information to the VISp than to MOp.

Visual responses of VISp neurons change in relation to the behavioral state of the animal^26^, and top-down projections are known to carry non-visual information to sensory areas^27,28^. By measuring pupil diameter and locomotor activity of the mice, each trial / stimuli presentation was assigned one out of three different behavioral states: aroused (>80^th^ percentile arousal events / pupil size per recording session, regardless if the animal was running), running (>80^th^ percentile running speed, per recording session), and neutral states (stationary, <20^th^ percentile arousal events / pupil size) (see Methods for details). Comparing the amplitude of the mean response to the preferred stimulus/stimuli of ACA and ORB axons in these states revealed that PFC axonal activity was also modulated by behavioral state. The normalized amplitude of visual responses was higher in an aroused and running state, in comparison to a neutral state (**Figure 2K**, and **Supplementary Figure 3E-F**). Notably, ORB axons displayed a more prominent change in response amplitude to visual stimuli between the neutral and aroused state. Their mean response amplitude in the neutral state was lower than that of ACA axons and higher than that of ACA axons in the aroused state (Gratings: neutral: p-value <0.001, aroused: p-value <0.001, running: p-value >0.05. Movies: neutral: p-value <0.001, aroused: p-value <0.001, running: p-value >0.05, one-way ANOVA with post-hoc Tukey test for multiple comparisons of means). The amplitude change in visual responses was higher for ORB axons, in comparison to ACA axons, during high arousal trials, i.e., the defined aroused state. However, this was not the case during trials with a lower arousal level (>40^th^ and <80^th^ percentile arousal events / pupil size, per recording session). Here, ACA axon activity response amplitude scaled more continuously with the level of arousal, while ORB axonal activity response amplitude instead increased sharply during high arousal trials only (gratings: neutral <40^th^ percentile: p-value >0.05, >40^th^ - <80^th^ percentile: p-value <0.001, >80^th^ percentile: p-value = 0.0061, one-way ANOVA with post-hoc Tukey test for multiple comparisons of means) (**Figure 2L**).

To summarize, ACA axon visual response amplitudes were more reflective of both the contrast of the visual stimuli and level of arousal, suggesting a modulatory effect onto VISp activity incorporating both visual stimulus information and a continuum of arousal state. ORB axon visual responses did not discriminate direction and contrast as well, but instead reflected a discrete behavioral state of high arousal, suggesting a more behavior-dependent role in modulating VISp activity. Comparing ACA and ORB input to VISp versus MOp revealed that while discrete traits such as contrast encoding and behavioral state modulation of visual responses were still preserved in ACA and ORB axon activity within MOp, the population-level strength of visual information was weaker in PFC projections to MOp. Thus, PFC regions preferentially send visual information to VISp compared to MOp.

### Visual information in ACA axons scales with arousal level

To directly assess whether the representation of visual stimuli in PFC axon population activity was influenced by behavioral state, we trained an SVM on all trials and evaluated the decoding accuracy of single-trial activity based on each trial’s behavioral state. Comparing decoding accuracy across different behavioral states revealed that ACA axon activity in VISp encoded visual stimulus identity significantly better during running and arousal trials compared to neutral trials (2-Way ANCOVA, ACA, running vs. neutral: p-value = 5.42e-27, arousal vs. neutral: p-value = 1.13e-23) (**Figure 3A-B**). Similarly, ORB axon population activity also decoded visual stimulus direction significantly better during running and arousal trials compared to neutral trials, though the improvement was smaller than in the ACA (2-Way ANCOVA, ORB, running vs. neutral: p-value = 2.14e-07, arousal vs. neutral: p-value = 3.04e-05) (**Figure 3C-D**). To further examine whether improvements in decoding accuracy were related to continuous arousal levels, similar to the ACA axon response amplitude to visual stimuli (**Figure 2L**), we analyzed the relationship between decoding accuracy and arousal level on a trial-by-trial basis. Plotting decoding accuracy for the largest subset of axons per trial against z-scored pupil size revealed a linear relationship between ACA axon decoding accuracy and pupil size (**Figure 3E**). Interestingly, this linear relationship was significant only for low-contrast stimuli (4% and 16%) but not for high-contrast stimuli (64%) (Ordinary Least Squares (OLS) regression, contrast 4%: p-value = 0.002836 slope = 0.468734, contrast 16%: p-value = 0.002320 slope = 0.422409, contrast 64%: p-value = 0.363367 slope = 0.109846). In contrast, ORB axon population activity showed no significant correlation between decoding accuracy of stimulus direction and pupil size at any stimulus contrast (contrast: 4% p-value=0.148045 slope=2.197141, contrast: 16% p-value=0.795380 slope=0.348075, contrast: 64% p-value=0.179783 slope=1.934175. (**Figure 3F**).

**Figure 3.**
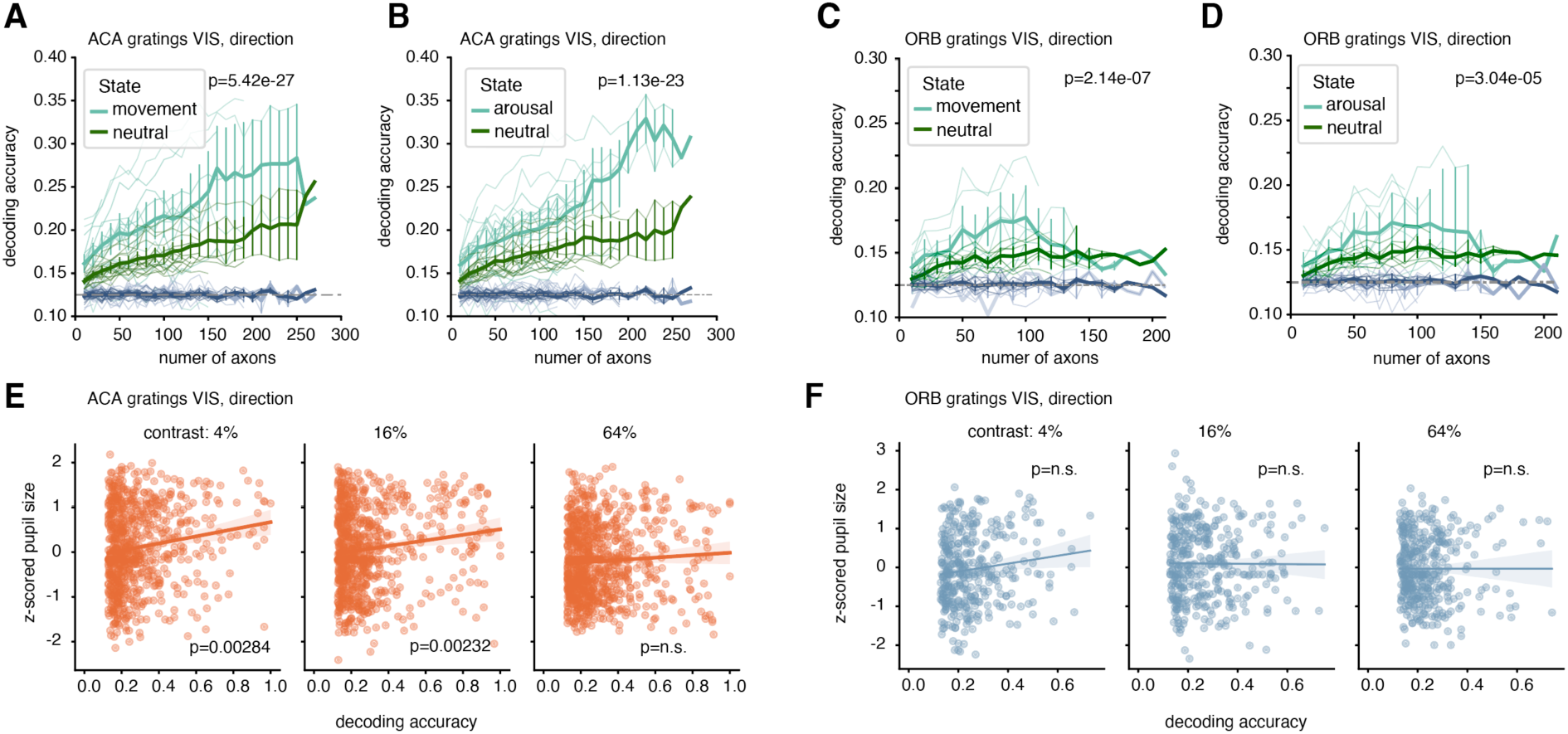
Decoding accuracy of visual information scales with arousal level in ACA axons. **(A)** Decoding accuracy of SVM decoder of drifting gratings directions trained on ACA axonal responses to drifting gratings (mean ΔF/F during stimuli on-time) in the VIS, and tested trial-by-trial. Singe trial decoding accuracy, averaged per session, is plotted for movement trials (light green) and neutral (dark green) trials. p-value from the variability in accuracy explained by group (movement vs. neutral). Direction: p-value: 5.42e-27. 18 fields of view across 4 mice. 855 running trials, 3420 neutral trials. **(B)** Decoding accuracy of SVM decoder of drifting gratings directions trained on ACA axonal responses to drifting gratings (mean ΔF/F during stimuli on-time) in the VIS, and tested trial-by-trial. Singe trial decoding accuracy, averaged per session, is plotted for arousal trials (light green) and neutral (dark green) trials. p-value from the variability in accuracy explained by group (pupil vs. neutral). Direction: p-value: 1.13e-23. 18 fields of view across 4 mice. 1036 arousal trials, 3455 neutral trials. **(C)** Decoding accuracy of SVM decoder of drifting gratings directions trained on ORB axonal responses to drifting gratings (mean ΔF/F during stimuli on-time) in the VIS, and tested trial-by-trial. Singe trial decoding accuracy, averaged per session, is plotted for movement trials (light green) and neutral (dark green) trials. p-value from the variability in accuracy explained by group (movement vs. neutral). Direction: p-value: 2.14e-07. 9 fields of view across 4 mice. 432 running trials, 1728 neutral trials. **(D)** Decoding accuracy of SVM decoder of drifting gratings directions trained on ORB axonal responses to drifting gratings (mean ΔF/F during stimuli on-time) in the VIS, and tested trial-by-trial. Singe trial decoding accuracy, averaged per session, is plotted for arousal trials (light green) and neutral (dark green) trials. p-value from the variability in accuracy explained by group (pupil vs. neutral). Direction: p-value: 3.04e-05. 9 fields of view across 4 mice. 432 arousal trials, 1728 neutral trials. Decoding accuracy of shuffled ACA/ORB axonal responses (movement/arousal, light gray; neutral, dark gray) plotted as comparison. Speed/Pupil value per trial is averaged value -1 sec before, during and after visual stimuli onset. Behavioral state was classified as the >80^th^ percentile trials per session (running speed: movement, pupil size: arousal, <80^th^ percentile events: neutral). Thick solid line: mean accuracy across sessions, error bars: 95% confidence interval (omitted when only one session contributed to a bin), thin lines: individual sessions, dashed line: chance level. p-value from 2-way ANCOVA to examine the effects of subset (number of axons) and behavioral state on the accuracy of decoding. The model included interaction terms to explore if the effect of subset on accuracy varies by group and data/shuffled data. **(E)** Linear regression lines fit to z-scored pupil size and decoding accuracy per trial, per contrast, of ACA axons to drifting gratings. The slopes and significance levels are determined using Ordinary Least Squares (OLS) regression. Single trials are represented by dots, shaded are 95% confidence interval. Number of trials per contrast: 4% n=710, 16% n=757, 64% n=834. p-value and slope per contrast: 4% slope-0.468734 p-value-0.002836, 16% slope-0.422409 p-value-0.002320, 64% slope-0.422409 p-value-0.002320. **(F)** Linear regression lines fit to z-scored pupil size and decoding accuracy per trial, per contrast, of ORB axons to drifting gratings. The slopes and significance levels are determined using Ordinary Least Squares (OLS) regression. Single trials are represented by dots, shaded are 95% confidence interval. Number of trials per contrast: 4% n= 352, 16% n= 348, 64% n= 364. p-value and slope per contrast: 4% slope-2.197141 p-value-0.148045, 16% slope-0.348075 p-value-0.795380, 64% slope-1.934175 p-value-0.179783.

These results suggest that ACA axons provide the VISp with a stronger representation of low evidence (low contrast) visual stimuli during high levels of arousal, whereas ORB axons’ representation of visual stimuli is strengthened during high levels of arousal but without a continuous modulation related to arousal level.

### ACA and ORB axonal activity reflects behavioral state differentially in VISp and Mop

Since PFC axons exhibited behavioral state-dependent modulation of response amplitude and population encoding of visual stimuli (**Figure 2**), we next addressed how ACA and ORB axons represent behavioral variables on a population level. In general, an overwhelming majority of both ACA and ORB axons showed a significant correlation of activity to pupil diameter, face movement, and locomotion in both the VISp and MOp (**Supplementary Figure 4A-D**). As ACA and ORB visual responses seems to scale differently with pupil dilation, we also evaluated whether ACA and ORB axonal population activity related better to the discrete velocity of movement (running speed or face velocity) and pupil size or to a binary measure of movement or arousal state and, in addition, if the axonal activity was differently modulated within the VISp and MOp (**Figure 4A-B**, and **Supplementary Figure 4A-D**). Comparing the strength of correlation (Spearman’s correlation Rho for axons with p-value <0.01 to both discrete and binary) for each axon to either the discrete or binary behavioral value revealed that both ACA and ORB axonal activity in the VISp had a stronger correlation to the binary movement value, while PFC axonal activity in the MOp was more correlated to the discrete running speed (**Figure 4B, left**). For pupil size and facial movement, ACA and ORB axons in both VISp and MOp correlated more strongly with discrete measures (pupil diameter and facial motion velocity) than with corresponding binarized values (dilated vs. non-dilated pupil; moving vs. still face) (**Figure 4B, right** and **Supplementary Figure 4A–D**).

**Figure 4.**
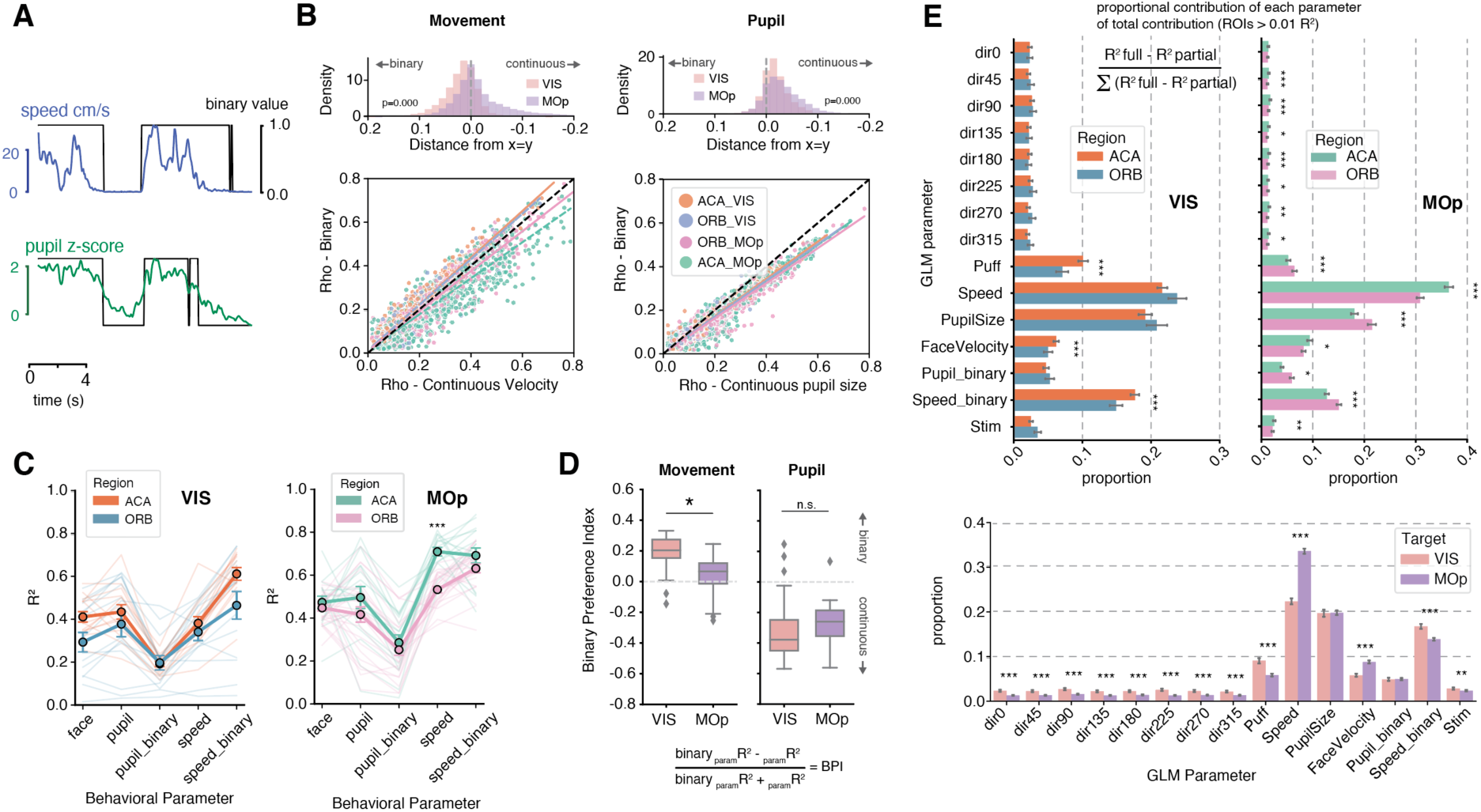
ACA and ORB axonal activity in the VISp is modulated by pupil size and a binary movement state. **(A)** Example trace of mouse movement on the running wheel and z-scored pupil size, overlayed with the binary state of movement (>0.5 cm/s) or pupil dilation (>1 z-score). **(B)** Down: Significant (p<0.01) Spearman correlation values (Rho) of each axon’s ΔF/F activity with the running speed (cm/s) against binary movement state (0 or 1) (left), or z-scored pupil size against binary dilated state (right). Each dot represents one axon color coded by the source and target area, only axons with significant correlation to both actual and binary value are included in the plot. Dashed lines are linear regression lines for each subset of axons relationship between the actual and binary movement or pupil size. Up: Distribution of absolute deviations from the identity line (|Δ| = Rho actual − Rho binary) for each group, summarizing how closely each axon’s correlations to actual versus binary measures align. Positive values indicate stronger correlation with the binary measure, while negative values indicate stronger correlation with the continuous measure. Differences between groups were tested using the Mann–Whitney U test, with p-values corrected for multiple comparisons using the Benjamini–Hochberg false discovery rate (FDR) method. Movement; ACA-VIS 2720 axons from 4 mice, ORB-VIS 1289 axons from 4 mice, ACA-MOp 4691 axons from 4 mice, ORB-MOp 4704 axons from 4 mice. Pupil; ACA-VIS 2573 axons from 4 mice, ORB-VIS 1191 axons from 4 mice, ACA-MOp 4299 axons from 4 mice, ORB-MOp 4494 axons from 4 mice. **(C)** Variance explained (R2) of behavioral parameters from a linear model (face (SVM z-scored), pupil (diameter z-scored), pupil binary value (0 or 1 if z-score > top 20^th^ percentile for that imaging session), speed (cm/s), speed binary value (0 or 1 if cm/s > 0.5) predicted by the axonal activity of ACA or ORB axons in the VIS (left) or MOp (right) during gratings sessions. Differences tested using the Mann–Whitney U test and Benjamini–Hochberg FDR correction; significance (* p < 0.05, ** p < 0.01, *** p < 0.001) is indicated above the bars. Dots: mean R2 of all imaging sessions, error bars: SEM, lines: individual sessions. Imaging sessions included are n=14 ACA-VIS, n=14 ORB-VIS, n=16 ACA-MOp, n=25 ORB-MOp. **(D)** Strength of correlation to actual vs binary value of movement (left) or pupil diameter (right) calculated as a binary preference index (BPI) by the variance explained for the binary versus actual value per recording session. BPI is plotted for each subset of axons based on target area. Each boxplot represents the quartiles of values, while the whiskers extend to show the rest of the BPI values, per imaging session, outliers represented as diamonds. * = p-value <0.05, one-way ANOVA with post-hoc Tukey test for multiple comparisons of means. **(E)** Variance explained (R2) of each parameter added to a linear model predicting the activity of single axons that has at least 1% activity explained by the model. Contribution of each parameter plotted as the proportional contribution of the total contribution explained per axon. Bars: mean proportion, error bars: 95% confidence interval. ACA-VIS 2940 axons, 22 imaging sessions, 4 mice. ORB-VIS 1444 axons, 19 imaging session, 4 mice. ORB-MOp 5015 axons, 28 imaging sessions, 4 mice. ACA-MOp 4999 axons, 21 imaging sessions, 4 mice. Down: An additional grouping compared VIS versus MOp (pooled across source areas), with differences tested using the Mann–Whitney U test and Benjamini–Hochberg FDR correction; significance (* p < 0.05, ** p < 0.01, *** p < 0.001) is indicated above the bars.

To further investigate to what extent PFC axonal activity tracked behavioral states, a generalized linear model (GLM) was fit to predict the fluctuation of each behavioral variable utilizing axonal population activity (see methods). PFC axonal activity predicted a large proportion of the observed behavior of the mice (face movement, pupil size, binary pupil size, speed, binary speed). Comparing the variance explained (R^2^) of each behavioral variable predicted by the model, PFC axonal activity in VISp was again observed to better predict a binary movement state rather than the speed of the animal (binary preference index, ACA-VIS, t-statistic: 8.65, p-value = 9.38e-07, ORB-VIS, t-statistic: 4.76, p-value = 0.00037, t-test of significant difference from 0). This was, however, not the case for pupil size where the prediction for actual pupil size outperformed the prediction of binarized pupil size (binary preference index, ACA-VIS, t-statistic: -15.457, p-value = 9.528e-10, ORB-VIS, t-statistic: -3.433, p-value = 0.004, t-test of significant difference from 0) (**Figure 4C-D**, and **Supplementary Figure 4E-f**).

Finally, a linear model was used to predict the activity of single axons based on the visual stimuli presented and the behavioral variables mapped during each imaging session (**Figure 4E**, and **Supplementary Figure 4G**). Comparing the proportional contribution of each parameter, to the total contribution of all parameters (for axons with a variance explained of R^2^>0.01), revealed that the most profound differences in how behavioral variables aid the prediction of PFC axonal activity was found not between the two source areas, the ACA and the ORB, but rather between the target regions, the VISp and the MOp. The proportional contribution of visual stimuli information to predict ACA and ORB axonal activity in the VISp was larger than in the MOp. Despite this, the behavioral parameters were more predictive of axonal activity than the visual stimuli in both the VISp and MOp. The animal’s speed was the largest proportional contributor to the variance explained for PFC axonal activity in the MOp, while the binary speed and pupil size were more prominent contributors to PFC-VISp activity in comparison to PFC-MOp (**Figure 4E, Supplementary Figure 4G**).

In summary, our analysis reveals a previously unrecognized functional organization of PFC axonal activity, encompassing both shared and unique activity profiles within PFC subregions and between target regions. This activity is characterized by four distinct traits: 1) ACA axons modulate VISp activity in response to both visual contrast and graded arousal level, 2) ORB axons reflect a more binary arousal-dependent modulation of VISp, 3) ACA and ORB projections to MOp preserve contrast encoding and arousal modulation, but with weaker visual information strength compared to VISp, and 4) PFC axonal activity in both VISp and MOp scales with pupil size, reflecting differential encoding of movement and arousal states, with movement modulation being binary in VISp and continuous in MOp (**Supplementary Figure 9**). Thus, while both the ACA and ORB preferentially convey visual information to VISp, the nature of information from ACA is finer-grained with respect to both visual contrast and arousal.

### ACA and ORB differentially modulate VISp visual responses

The next pivotal question was how the axonal activity observed in our experiments above influences the activity of VISp neurons, and importantly the representation of visual stimuli. VISp activity has been shown to be altered by optogenetic activation of ACA axons in the VISp, which induce a sharpening of the tuning curve of visually responsive excitatory neurons via selective activation of local interneurons^17^. How endogenous, behaviorally driven PFC activity influences the encoding of visual information in the visual cortex, and in particular how discrete subregions of the PFC could participate in this process, has not previously been investigated. To address this, 2-photon imaging of VISp somatic activity was performed in both the presence and absence of ACA or ORB input. retroAAV-Cre was injected into the VISp, followed by AAV-DIO-hM4D(Gi) injected into the ACA or the ORB, allowing for projection specific inhibition of ACA or ORB neurons targeting VISp. A subsequent injection of AAV-hSyn-GCaMP7f in the VISp allowed for simultaneous imaging of VISp neurons with or without ACA or ORB feedback (**Figure 5A-B**). The same neurons were imaged across two days, in which the mouse received an injection of saline on one day and CNO (Clozapine-N-oxide, 1mg/kg) on the other (**Figure 5C**). Recorded neurons were then mapped across days (see Methods) to ensure that the neuronal activity recorded in the presence and absence of PFC modulation could be evaluated on a single-neuron level (dataset referred to as ‘matched’ throughout the paper). Awake behaving mice were free to run on a running-wheel as before, and VISp neuronal responses were recorded with 2-photon imaging (**Figure 2D**, **Figure 5D**). Visually responsive neurons were identified to an equal extent in the CNO and saline condition and the behavior of the mice (face movements, pupil size, running speed) was not significantly changed (Students paired t-test, p-value >0.05) (**Figure 5E**, and **Supplementary Figure 5**).

**Figure 5.**
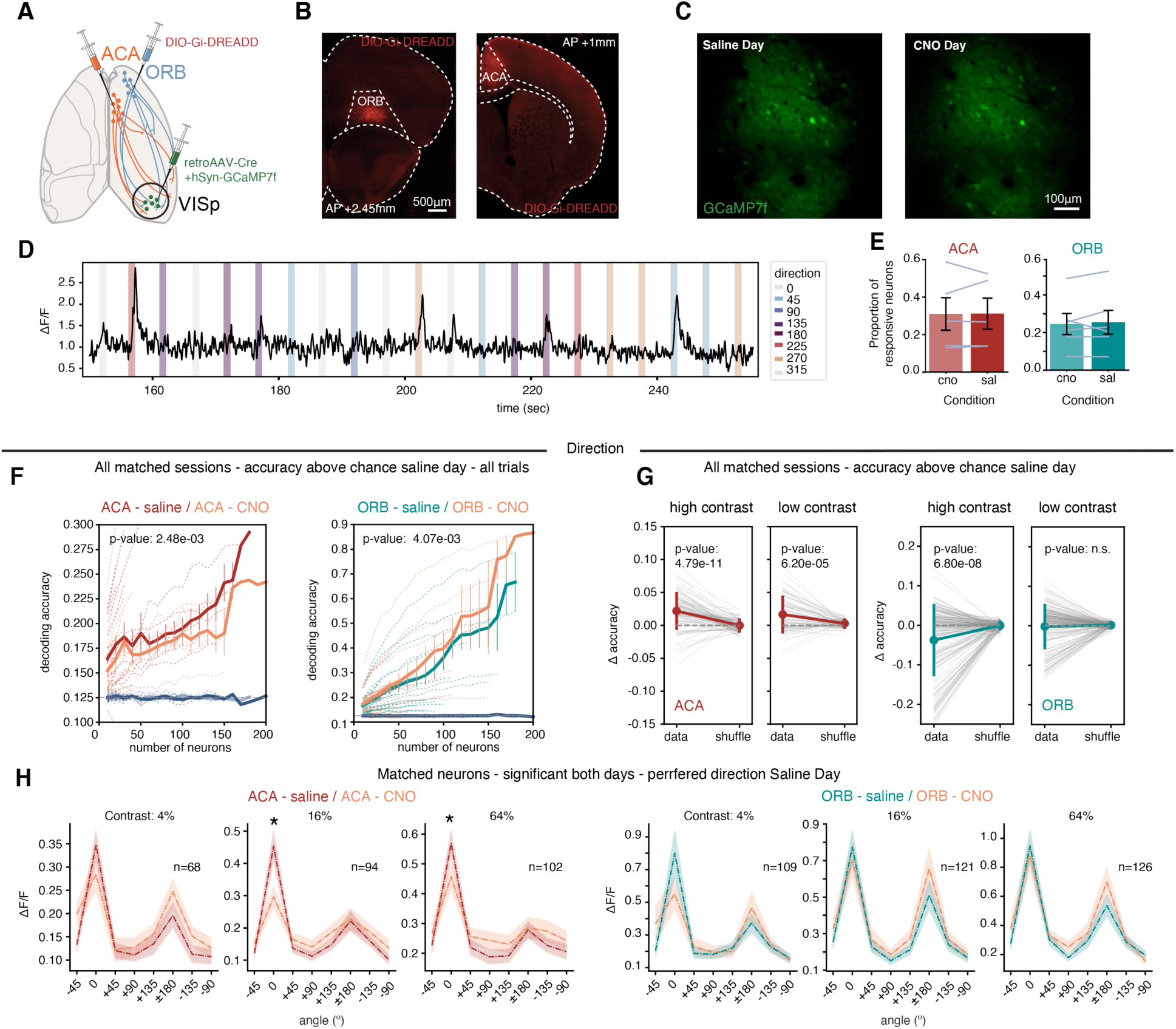
ACA and ORB differentially modulate visual responses of VISp neurons. **(A)** Experimental strategy. retroAAV-Cre and GCaMP7f (AAV-hSyn-DIO-GCaMP7f) was injected unilaterally into the VIS, followed by and injection of Cre-dependent inhibitory DREADDs (AAV-DOI-hM4D(Gi)-mCherry) into the ACA or the ORB. A 3mm cranial window was implanted above the VIS. **(B)** Representative images of viral injection sites in the ACA (left) and the ORB (right). Replicated in n=5 ACA, n=6 ORB mice. **(C)** Example field of view from 2-photon imaging of GCaMP7f expressing neurons in VIS across two days of imaging. **(D)** Activity trace (F) from one example VIS neuron, with the representation of drifting gratings color coded by the direction of grating displayed. **(E)** Proportion of visually responsive VIS somas in each experimental condition. Bars display mean of proportions, error bars: SEM, lines represent individual animals. Saline condition: 1760 neurons, CNO condition: 1832 neurons, across n=5 mice ACA. Saline condition: 3228 neurons, CNO condition: 2701 neurons, across n=6 ORB mice. **(F)** Decoding accuracy of SVM decoder of drifting gratings directions trained and tested on saline day (red-ACA, green-ORB) or CNO day (orange) VIS soma responses (mean ΔF/F during stimuli on-time, per imaging session and field of view. Decoding accuracy of shuffled soma responses saline day (light gray) or shuffled soma responses CNO day (dark gray) plotted for comparison. All paired recording sessions with a minimum accuracy above chance level on saline day included in the analysis. Thick solid line: mean accuracy across sessions, error bars: 95% confidence interval, thin lines: individual imaging sessions, dashed line: chance level. ACA: 26 paired recording sessions n=5 mice, ORB: 33 paired recording sessions n=6 mice. 2-way ANCOVA to examine the effects of subset (number of neurons) and experimental condition (saline/ CNO) on the accuracy of decoding. The model included interaction terms to explore if the effect of subset on accuracy varies by group and data/shuffled data. p-value from the variability in accuracy explained by group (saline vs. CNO). **(G)** Decoding accuracy of SVM decoder of drifting gratings directions trained and tested on VIS soma responses (mean ΔF/F during stimuli on-time) on either high contrast (64%) or low contrast (16%, 4%) stimuli. Change in decoding accuracy per paired recording session (field of view) and subset (number of neurons), comparing real and shuffled data. Mean Δ accuracy across sessions, error bars: standard deviation, grey lines are paired actual and shuffled data from same session and subset. p-value from paired t-test between real and shuffled data. High contrast trials; ACA: 20 paired recording sessions n=5 mice, ORB: 31 paired recording sessions n=6 mice. Low contrast trials; ACA: 20 paired recording sessions n=5 mice, ORB: 30 paired recording sessions n=6 mice. **(H)** Tuning curves of matched neurons that were significantly visually responsive on both days to the same direction(s), aligned to their preferred direction, plotted per contrast. Preferred direction was determined by the highest amplitude of response on the saline day. Thick line: mean value, shaded area: 95% confidence interval. Left; ACA-DREADDs 16% p-value: 0.0001, 64% p-value: 0.0001, paired t-test ΔF/F responses across matched neurons per direction. Left; ACA-DREADDs 4%: 68 neurons, 16%: 94 neurons, 64%: 102 neurons, from 5 mice. Right; ORB-DREADDs 4%: 109 neurons, 16%: 121 neurons, 64%: 126 neurons, from 6 mice.

To evaluate the accuracy with which visual stimuli could be decoded from VISp activity in the presence and absence of PFC feedback, a SVM was trained and tested on the mean ΔF/F activity of VISp neurons during stimulus presentation (stratified by contrast and direction), on each recording session separately. Each pair of recording sessions (same field of view on saline day and CNO day) were included if the minimal decoding accuracy was above chance on the saline day, and the decoding accuracy was compared across each paired recording sessions. Intriguingly, ACA and ORB modulation of VISp activity had opposite effects on the accuracy with which the direction of drifting gratings could be decoded: removing ACA input to the VISp decreased the decoding accuracy, while removing ORB input improved the decoding accuracy of visual stimuli (2-Way ANCOVA, gratings: ACA p-value = 0.0025, ORB p-value = 0.0041) (**Figure 5F**). While averaged decoding accuracy was higher in the ORB-saline group (**Figure 5F**), this likely reflects session variability, as the direction and significance of modulation effects were consistent across paired sessions and different type of stimuli (**Figure 5G and Supplementary Figure 6**). Given that ACA axons in VISp had a higher sensitivity to contrast in comparison to ORB axons, the effect of contrast on the decoding accuracy was evaluated by training a SVM on trials with either high (64%) or low (4% and 16%) contrast stimuli. The removal of ACA input worsened the decoding accuracy of both high and low contrast gratings (paired t-test, high contrast p-value = 4.79e-11, low contrast p-value = 6.20e-05) (**Figure 5G, left**). The effect of contrast was more pronounced with regard to the decoding accuracy of natural movies with and without ACA input (paired t-test, high contrast p-value = 0.22, low contrast p-value = 1.58e-12) (**Supplementary Figure 6C-D**). Interestingly, removing ORB input improved the decoding accuracy of both high contrast gratings (paired t-test, high contrast p-value = 6.80e-08, low contrast p-value = 0.37) as well as natural movies (paired t-test, high contrast p-value = 1.3e-5, low contrast p-value = 0.85) (**Figure 5G, right,** and **Supplementary Figure 6A-B**). Investigating the visual tuning of individual VISp neurons at different contrasts also revealed that ACA modulation affected the response amplitude of visually responsive neurons at their preferred direction at multiple contrasts, while the removal of ORB modulation had no effect (paired t-test, p-value <0.00625) (**Figure 5H**).

To summarize, discrete PFC subregions differentially influence the responses of VISp neurons to visual stimuli. ACA input enhances the directional selectivity of VISp neurons as well as stimulus encoding (**Figure 5F and H, left**), while conversely ORB input appears disruptive to the encoding of high-contrast stimuli without impacting the directional tuning of individual neurons (**Figure 5F and H, right**).

### ACA modulation of VISp is dependent on arousal state

Behavioral state changes are known to modulate cortical activity, and changes in VISp activity in particular have been linked to discrete behavioral variables such as body movement, pupil size, and face movements^26,29^. Since ACA and ORB axonal activity were modulated by behavioral variables, we next questioned whether their modulation of VISp activity is related to different behavioral states. As described previously^26,30^, visually responsive VISp neurons increased their average and peak ΔF/F response amplitude to visual stimuli during a state of arousal or of running in comparison to a neutral state (**Figure 6A-B, saline condition**). Removing either ACA or ORB input caused the observed increase in response amplitude in the aroused state to decrease (p-value <0.01, one-way ANOVA with post-hoc Tukey test for multiple comparisons of means) (**Figure 6A-B**). Furthermore, removing ORB modulation caused the observed increase in response amplitude in the running state to decrease (p-value <0.01, one-way ANOVA with post-hoc Tukey test for multiple comparisons of means) (**Figure 6A-B**). Importantly, this was not a consequence of altered arousal levels or overall movement, as the mice’s behavior (face movements, pupil size, running speed) was not significantly changed between saline and CNO conditions (Students paired t-test, p-value>0.05) (**Figure 5E**, and **Supplementary Figure 5**). In addition, the latency of visual responses was not affected by ACA or ORB modulation independent of state (**Supplementary Figure 7A**).

**Figure 6.**
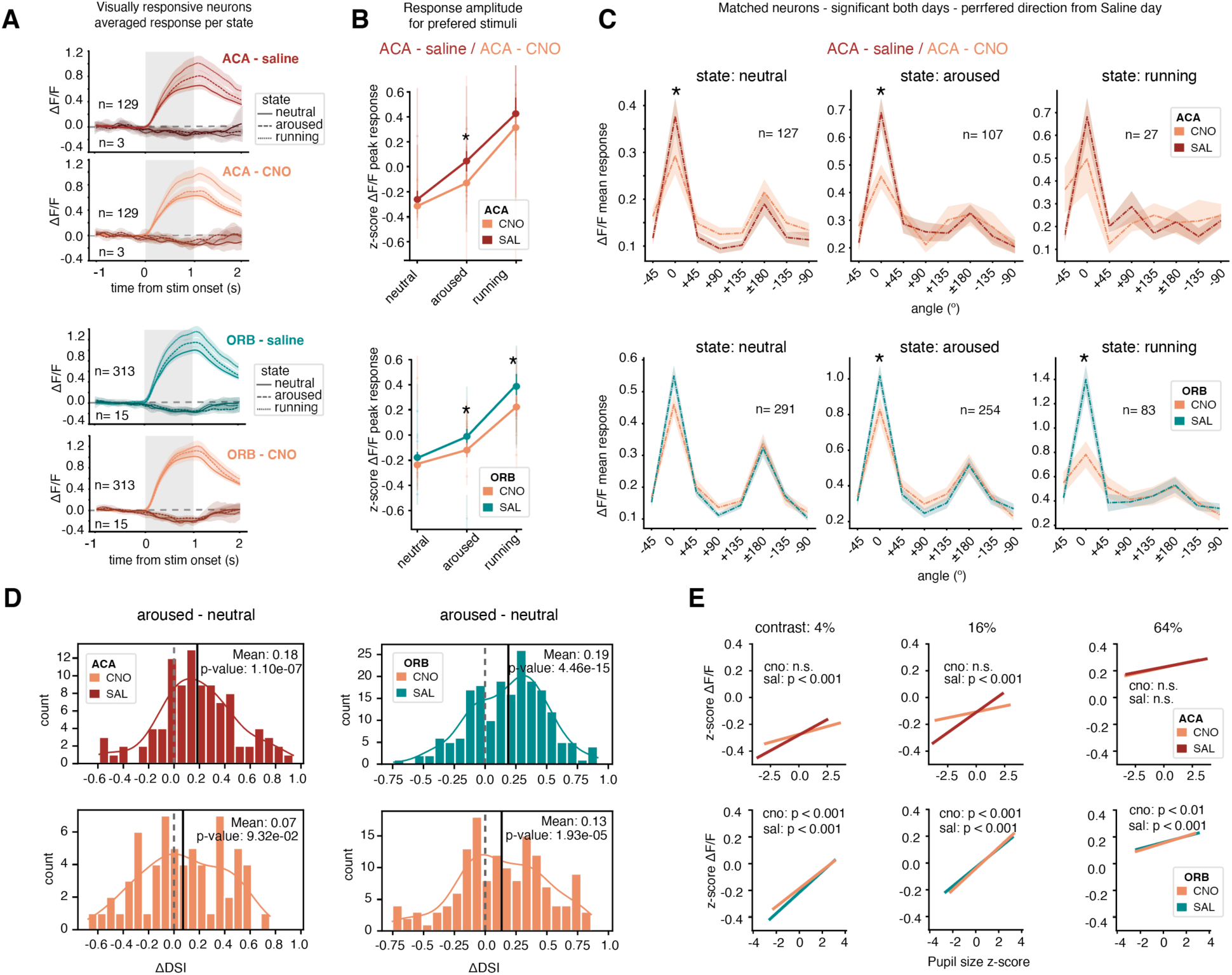
ACA modulation of responses to weak stimuli relates to arousal state. **(A)** Averaged population responses of matched neurons that were significantly visually responsive on both days to the same direction(s) plotted per behavioral state, and response direction. Trials/responses were classified as belonging to a certain behavioral state as the top 20 percentile events per session (running speed: movement, pupil size: arousal, less than top 20 percentile events: neutral). VISp neurons were classified as positively or negatively tuned based on whether their mean response during stimulus presentation was above or below zero. Top; ACA-DREADDs 132 neurons from 5 mice, bottom; ORB-DREADDs 328 neurons from 6 mice. Thick line: mean value, shaded area: 95% confidence interval. **(B)** Standardized ΔF/F peak responses during stimuli on-time of matched neurons that were significantly visually responsive on both days to the same direction(s), plotted per behavioral state. Behavioral state was classified as in A. ΔF/F mean responses are z-scored across each neuron. Bars: mean standardized ΔF/F mean responses across axons, error bars: 95% confidence interval, dots: individual animals. * p-value < 0.01, one-way ANOVA with post-hoc Tukey test for multiple comparisons of means. Top; ACA-DREADDs 132 neurons from 5 mice, bottom; ORB-DREADDs 328 neurons from 6 mice. **(C)** Tuning curves of matched neurons that were significantly visually responsive on both days to the same direction(s), aligned to their preferred direction, plotted per behavioral state. Preferred direction was determined by the highest amplitude of response on the saline day, and a neuron was only included if at least one trial per direction existed for that state. Thick line: mean value, shaded area: 95% confidence interval. Top; ACA-DREADDs neutral state p-value: <0.0001, aroused state p-value: <0.0001. Bottom; ORB-DREADDs aroused state p-value: <0.0001, running state p-value: <0.0001, paired t-test ΔF/F responses across matched neurons per direction. Top; ACA-DREADDs neutral state: 127 neurons, arousal state: 107 neurons, running state: 27 neurons, from 5 mice. Bottom; ORB-DREADDs neutral state: 291 neurons, arousal state: 254 neurons, running state: 83 neurons, from 6 mice. **(D)** Histogram of values of change of Direction Selectivity Index (ΔDSI) from a neutral state to an aroused state (top 20 percentile pupil events). ΔDSI values stem from matched neurons that were significantly visually responsive on both days to the same direction(s), aligned to the preferred direction on the saline day. Dashed line indicates 0, and the solid line is the mean of ΔDSI values plotted. P-value from one-sampled t-test against 0. ACA-DREADDs saline: 97 neurons CNO: 64 neurons from 5 mice. ORB-DREADDs saline: 146 neurons CNO: 191 neurons from 6 mice. **(E)** Linear regression lines fit to z-scored ΔF/F single trial activity against z-scored pupil size, per condition (sal/cno) and contrast. Single trial activity is included from matched neurons that were significantly visually responsive on both days to the same direction(s). ΔF/F mean responses for stimuli on-time per trial are z-scored across each neuron and condition. The slopes and significance levels are determined using Ordinary Least Squares (OLS) regression. Top: ACA-DREADDs 152 neurons from 5 mice, total of 49290 trials, 24737 CNO trials and 24553 saline trials. Bottom: ORB-DREADDs 380 neurons, total of 120652 trials, 60336 CNO trials and 60316 saline trials.

Further investigating the effect of ACA and ORB modulation on VISp neuron tuning curves revealed that the significant drop in response amplitude between saline and CNO days was due to a drop in the average response amplitude to the preferred direction of drifting gratings, and not simply all, or similar, orientations (**Figure 6C**). Comparing the mean ΔF/F response amplitude of matched VISp neurons aligned to their preferred direction on the saline day showed that the response amplitude to the preferred direction decreased in the neutral and aroused state in the absence of ACA modulation, while the responses to other directions remained unchanged (p-value <0.00625, paired t-test). In comparison, while the removal of ORB modulation yielded a similar change in mean amplitude of response to the VISp neurons’ preferred direction of stimuli during aroused and running states, this was not observed in the neutral state (p-value <0.00625, paired t-test) (**Figure 6C**).

Given that the effect of removing PFC input appeared specific to the preferred direction, we next investigated how the direction selectivity index (DSI) of VISp neurons changed between behavioral states in the presence or absence of ACA or ORB input (**Figure 6D** and **Supplementary Figure7A**). The analysis revealed that in the absence of ACA input, the increase in DSI values seen during a state shift from neutral to aroused was lost (one-sampled t-test against 0, saline: p-value = 1.10e-7 mean = 0.18, CNO: p-value = 0.093 mean = 0.07) (**Figure 6D, left**). In contrast, the removal of ORB modulation only resulted in a small drop in the increase in mean DSI values from the neutral to aroused state, and there was still a significant increase in DSI values due to the state shift in the absence of ORB modulation (one-sampled t-test against 0, saline: p-value = 4.46e-15 mean = 0.19, CNO p-value = 1.93e-5 mean = 0.13) (**Figure 6D, right**).

Finally, we addressed how the change in response amplitudes during a state shift from neutral to an aroused state related to the contrast of the stimuli, since we had found a differential effect of ACA and ORB modulation on VISp neurons’ encoding of stimuli of different contrasts (**Figure 5G**). Interestingly, ACA modulation of low-contrast stimuli was highly dependent on the discrete arousal state (pupil size) of the animal, i.e., a strong correlation of amplitude of response and pupil size, that was lost in the absence of ACA modulation (Ordinary Least Squares (OLS) regression, Saline: contrast 4%: p-value = 4.87e-5 slope = 0.05, contrast 16%: p-value = 3.72e-8 slope = 0.06, contrast 64%: p-value = 0.16 slope = 0.01, 24553 trials, CNO: contrast 4%: p-value = 0.043 slope = 0.02, contrast 16%: p-value = 0.1 slope = 0.02, contrast 64%: p-value = 0.10 slope = 0.02, 24737 trials, n= 152 neurons) (**Figure 6E**). The modulation of response amplitude of low-contrast stimuli in relation to arousal state was unaffected by the removal of ORB modulation onto VISp neurons (Ordinary Least Squares (OLS) regression, Saline: contrast 4%: p-value = 4.5e-21 slope = 0.07, contrast 16%: p-value = 2.61e-27 slope = 0.08, contrast 64%: p-value = 4.3e-4 slope =0.03, 60316 trials, CNO: contrast 4%: p-value = 6.8e-25 slope = 0.08, contrast 16%: p-value = 8.3e-22 slope = 0.07, contrast 64%: p-value = 0.001 slope = 0.02, 60336 trials, n= 380 neurons). Examining the change of visual responses during a state shift from a neutral to a running state revealed an increase in DSI values, and a relationship between the running speed of the animal and increase in amplitude of visual responses, but this movement related modulation was unaffected by the removal of ACA input, and only moderately affected by ORB modulation (**Supplementary Figure 7B-C**). The observed scaling of the amplitude of response to visual stimuli with the speed of running is likely to be influenced by neuromodulatory or locomotion centers in the brain, and not the PFC, as ACA and ORB axonal activity was observed to reflect a binary movement state in the VISp.

A GLM was used to evaluate how all visual and behavioral parameters contributed to the prediction of VISp neurons’ activity, in the presence and absence of ACA or ORB modulation. Evaluating the proportional contribution of each parameter to the variance explained by the model revealed no major differences in how each parameter contributed to the prediction of activity with or without ACA or ORB modulation (**Supplementary Figure 8A**). These results suggests that although the ACA and ORB modulate the pattern of activity and visual responses of VISp neurons, these modulatory inputs do not alter the proportional representation of sensory and behavioral information in VISp neurons.

To conclude, our data indicates that ACA modulation strengthens the representation of visual stimuli, particularly low contrast (low evidence) stimuli, by increasing the response amplitude of VISp neurons to their preferred direction, resulting in an increase in decoding accuracy. The modulation of visually responsive VISp neurons by the ACA is dependent on arousal, and correspondingly the increase in response amplitude to low-contrast stimuli seen in VISp neurons is the most profound during high arousal states. ORB modulation also contributes to the increase in VISp neuron response amplitudes towards the preferred direction, especially during running and high arousal states.

## Discussion

In this study, we have demonstrated that the VISp and MOp receive direct axonal projections from two discrete PFC subregions, the ACA and the ORB. The two projection populations target excitatory as well as all inhibitory cell types, suggesting a rich influence on microcircuits in VISp and MOp. Indeed, the ACA and ORB have both shared and unique modulatory influences on VISp neuronal activity. The two projection populations showed distinct differences in their encoding of visual stimuli, whereby ACA axons were more visually responsive and scaled their response with the contrast of stimuli while ORB axons did not. Both PFC projections were modulated by behavioral state, consistent with the finding that both ACA and ORB neurons are influenced by arousal and movement, likely through neuromodulatory inputs^32,33^.

Comparing PFC axonal activity between VISp and MOp revealed that PFC feedback to VISp carries a stronger representation of visual information at the population level, while feedback to MOp is more strongly correlated with the animal’s speed, with body movement being the most prominent predictor of PFC axon activity in MOp. Notably, the two PFC regions diverged in their representation of body movement in downstream regions: in VISp, movement was encoded as a binary state, while in MOp, movement modulation was continuous, relating the axonal activity to discrete velocities (Supplementary Figure 9).

Further comparisons of ACA and ORB axon activity in superficial layers of both VISp and MOp revealed differences in visual encoding, such as contrast, which were preserved across both target regions. ACA and ORB axon activity were both modulated by arousal level, but while ACA axonal activity was graded in response to arousal, ORB activity was binary, modulated only at high levels of arousal. Therefore, both the ACA and the ORB likely have distinct neuronal subpopulations targeting either both downstream regions or one more exclusively, supporting discrete functional subnetworks within ACA and ORB feedback projections which earlier have only been described on the anatomical level^8^. Removing ACA or ORB input to the VISp had differential effects on the VISp population encoding of visual stimuli. On a single-cell level, ACA modulation increased VISp neurons’ response amplitude towards their preferred direction. While ORB modulation had no effect on their tuning curves in a neutral state, ORB modulation increased the response amplitude to their preferred stimuli during a state of high arousal. Thus, our data indicates that the PFC subregions provide both shared and diverse feedback modulation to the VISp and broader cortex.

ACA and ORB projections were here shown to innervate different layers of the VISp, with ACA projections targeting layers 1 and 6, and ORB projections targeting layers 1 and 5 (**Figure 1D-F**). These PFC projections have previously been shown to target and activate different components of VISp microcircuitry^7,8,16^, which here was even further supported by our anatomical tracing and observations of their distinct influence on visual processing. Intriguingly, in this study, cortex-wide tracing of ACA and ORB axons revealed that this pattern of innervation, i.e., differential targeting of deep layers, was consistent across what is considered the ‘medial network’ of PFC axonal projections or ‘visual and medial network’ of cortex-wide connectivity ^5,7,8^. Not only did the ACA and ORB target different layers of the VISp, but this pattern was here shown to be consistent in all visual cortex regions and RSP (**Supplementary Figure 1F and H**). ACA and ORB axonal innervation of regions belonging to the ‘sensorimotor network’ or ‘central network’ showed no such differences, instead they shared a rather similar innervation of both deep and superficial layers (**Supplementary Figure 1F and H**). These findings are also in line with the presented ACA and ORB axonal activity imaging results, indicating differential representation and modulation of visual stimuli in VISp, but a similar representation of movement in MOp, suggesting discrete functional subnetworks of ACA and ORB projection populations targeting either the VISp or MOp. Future work is needed to examine the feedback from ACA and ORB to deeper cortical layers of VISp to see if the activity profiles reported here are distinct to the superficial input, as it is likely separate PFC neuronal populations targeting the deeper layers of the VISp^9^.

Visually responsive ACA axons were found at a similar proportion in our study (**Figure 2F-G**) to what has been reported previously in both ACA axons in VISp^27^ and ACA cell bodies ^33^. In addition, here the visual responses of ACA axons were shown to scale with the contrast of the stimuli in both the VISp and MOp (**Figure 2h-i and Supplementary Figure 2A-D**). Conversely, ORB axons responsive to visual stimuli were less prevalent in both the VISp and MOp, and visually responsive ORB axons did not scale their responses to the contrast of the visual stimuli to the same extent as ACA axons. The differential encoding of visual information in the two projections populations are likely due to differences in whole-brain connectivity. The ACA is known to predominantly share connectivity with the ‘medial network’ of the cortex, including visual cortex regions, RSP, and auditory cortex, in addition to visual thalamic nuclei and areas in the striatum with input from the aforementioned regions^7,34–36^. The ORB on the other hand, while also having connectivity with the medial network, is connected to the somatosensory and motor cortical and thalamic regions to a much larger extent than the ACA ^7,34–36^. Interestingly, the differences in visual responses, such as contrast sensitivity and behavioral state modulation, observed in ACA and ORB axons were consistent across the two downstream regions investigated (i.e., VISp and MOp), although the strength of population encoding for visual information differed (**Figure 2E**, and **Supplementary Figure 2**). This observation suggests that the visually responsive axons in the MOp could be collaterals of axons with a primary branch belonging to the medial network of the cortex, a projection profile which has been identified previously in both the ACA and the ORB^8^.

An important aspect of sensory responses identified in the mouse cortex is the influence of behavioral state on the representation of sensory information; in particular, how the representation of visual information changes based on the movement and arousal level of the animal^26,37^. In this study, visual responses of both ACA and ORB axons were found to also reflect the behavioral state of the animal (**Figure 2K-L**). As mentioned earlier, it is probable that state and movement-dependent inputs target both the ACA and ORB, in addition to other cortical regions, and that this modulation is therefore reflected in the activity of their axons^32,33^. In particular, the ACA axons’ response amplitude to visual stimuli, as well as, the decoding accuracy of ACA axon population activity, scaled with arousal level (pupil size) (**Figure 2L and 3E**). However, the visual responses of ORB axons had a higher increase in response amplitude, and decoding accuracy, selectively during high arousal states, but did not scale with arousal level (**Figure 2L and 3F**). Therefore, the incorporation of state-dependent modulation into the activity of these discrete projection populations appears distinct. The ACA and the ORB projections to VISp have previously been investigated in relation to visual learning and selective visual attention^17,18^, and the unconstrained viewing of visual stimuli in the present data reveals that natural state transitions impact ACA and ORB axonal activity and could subsequently influence the encoding of visual stimuli in the VISp.

The vast majority of ACA and ORB axons’ activity correlated to several of the behavioral parameters measured in the above experiments i.e., face movements, running speed, and pupil size **(Supplementary Figure 4A and C).** Intriguingly, the way in which ACA and ORB axons altered their activity in relation to discrete behaviors differed between axons in the VISp and MOp: while ACA and ORB axons’ activity in VISp would increase or decrease in relation to a binary stationary/moving state, the absolute speed of the animal held no additional explanatory power. This was not the case of ACA and ORB axons in the MOp, where instead the activity of ACA and ORB axons correlated more closely to the measured speed of the animal (**Figure 4A-D**), further highlighting that distinct movement-related neuronal subpopulations within ACA and ORB selectively target either the VISp or MOp. This is of interest since one can envisage a scenario in which PFC modulation of VISp neuronal activity is altered depending on the behavioral state of the animal, including how the visual scene is expected to change upon moving, but the discrete velocity of the animal’s movement is less relevant. In the MOp however, it would be more important for the PFC to provide modulation to MOp neurons that is related to the actual movements or velocity of the animal, as a central hub of motoric actions in the cortex. The type of binary movement state signal identified here in the PFC-VISp axons has also been identified previously in the activity of cholinergic axons from the basal forebrain projecting to the VISp, and this activity was suggested to relate to movements that result in changes to visual input^38^. Importantly, ACA axonal activity in the VISp may predict visual information to appear based on previous experiences in visual space^39^, and visual flow due to movement^28^. The activity of ACA and ORB axons we observe could serve a similar function, however these hypotheses cannot be fairly addressed in our current data.

Although axonal activity imaging provides a representation of what information is communicated to a downstream region, the theory of feedback modulation is founded on the ability of this input to change how the local circuitry processes or represents sensory information^3,4,40^. In this study, 2-photon imaging and chemogenetic interventions were used to observe how the same population of VISp neurons represented visual stimuli in the presence and absence of ACA or ORB feedback modulation. The absence of ORB modulation did not change the shape of averaged tuning curves of visually responsive VISp neurons, but it did increase the decoding accuracy of stimulus identity for high contrast stimuli on a population level (**Figure 5F-H**, and **Supplementary Figure 6**). The ORB has repeatedly been linked to reward and value related encoding and learning^41–44^, and has been directly implicated in modulating VISp responses by outcome expectation^18^. The drop in decoding accuracy of high contrast stimuli in the presence of ORB modulation was therefore of particular interest in relation to previous work that showed the necessity of the ORB–VIS circuit to reducing VISp responses towards non-relevant (non-rewarded) stimuli^18^. One hypothesis is that in a passive setting, the ORB would reduce the representation of prominent high contrast stimuli that is irrelevant, or unrewarded, in the current context. Another possibility is that ORB modulation related to movement interferes with visual cortex activity, similar to what has been described previously for movement signals in the primate visual cortex^45,46^.

In this study, the absence of ACA input decreased the average response amplitude of visually responsive neurons to their preferred stimulus. At the population level, this led to reduced decoding accuracy of stimulus identity, particularly for low-contrast stimuli (**Figure 5F-H, Supplementary Figure 6**). Previous work has shown that optogenetic activation of the ACA–VISp circuit enhances VISp somatic responses, especially for their preferred stimulus direction^17^. However, Zhang et. al. did not examine how ACA input modulates VISp responses across different stimulus contrasts. Furthermore, endogenously heightened ACA axonal activity in VISp does not reflect a similar sharpening of VISp neurons’ tuning curves^27^. Rather, endogenous ACA axonal activity levels observed in VISp, on a population level, are strongly related to motoric actions and behavioral state ^27,28,39,47^. Importantly, our findings unify these prior observations by showing that behavioral state modulates visual encoding by ACA axons in VISp, in particular the representation of low-contrast stimuli during arousal (**Figure 2 and 3**). Consistent with this, we also observed that ACA feedback modulation increased the response amplitude of VISp neurons towards their preferred stimuli, particularly at low contrasts, and that this modulation scaled with arousal level (**Figure 6D-E**). The ACA therefore likely works to enhance the representation of weak or uncertain stimuli during states of arousal, and likely during states of allocated attention. Our findings also reveal a similar dichotomy in ACA and ORB modulation during passive viewing to that observed during visually guided behavior in mice^17,18^ as well as a suggested functional dichotomy in the primate^49^.

To conclude, our study reveals that the PFC targets the VISp through two distinct populations of projection neurons originating from the ACA and the ORB. ACA and ORB axons have distinct laminar distributions in VISp reflecting their potentially complementary roles in the modulation of visual processing. ACA projections are more visually responsive and scale their response to the contrast of visual stimuli, while ORB projections do not exhibit such scaling. The differential encoding of visual stimuli by the two PFC subregions are likely consequences of brain-wide connectivity and differential access to sensory information. Both types of PFC projections are modulated by behavioral states, showing increased response amplitudes during high arousal and movement, indicating a shared movement and arousal-related activation of ACA and ORB projection neurons, albeit with distinct scaling. The distinct representation of visual information and differential encoding of movement in PFC axons within the MOp and the VISp is consistent with a mixture of shared and distinct projection sub-networks within each PFC subregion. The removal of each PFC subregion’s input to the VISp results in both unique and shared consequences on the activity of VISp neurons: neuronal activity modulated by high arousal and movement are both decreased, while the encoding of visual information worsens in the absence of ACA input and is surprisingly enhanced in the absence of ORB input (**Supplementary Figure 9**). Such a differential impact likely reflects distinct local microcircuitry that is targeted by ACA and ORB inputs to VISp, and unraveling the interplay of these downstream targets is a critical goal for future work.

## Acknowledgements

This research was supported by the Wenner-Gren foundations Postdoctoral Fellowship WGF2020-0019 and NIH grant 1K99EY035752 (SÄR), NIH grant F32EY032756 (KRJ), NIH grants R01MH133066 and R01NS130361, and The Picower Institute Innovation Fund (MS). We thank Yi Ning Leow and Greggory Heller for thoughtful discussions, Alexandria Barlowe and Jonathan Harpe for assistance with histology, Marco Celotto for aiding with the decoding analysis, and Cantin Ortiz for aiding with the creation of the axon tracing pipeline.

## Author contributions

SÄR conceived the study, designed and performed the experiments, analyzed the data and wrote the manuscript. YO designed the SVM decoder and Generalized Linear Model. KRJ built the behavioral rig, aided with experimental design and analysis, and provided extensive input to the manuscript. EO performed the immunohistological analysis of the ATLAS virus tracing. HH and DBA provided the unpublished ATLAS virus system and aided with its use. MS contributed to the direction of data analysis and the development of concepts presented, and provided thoughtful comments on the manuscript. All authors provided input to the manuscript.

## Competing interests

The authors declare no competing interests.

## METHODS

### Animals

All experiments were carried out in adult male and female mice under protocols conforming to NIH guidelines approved by the MIT Animal Care and Use Committee. Wild type C57BL/6J, Jackson stock no. 000664, or transgenic mouse line VIP-Flp: Vip^tm2.1(flpo)Zjh^/J, Jackson stock no. 028578 were used. All transgenic mice used in experiments were heterozygous for the transgenes. Mice were maintained under standard housing conditions with a 12-hour light cycle and with *ad libitum* access to food and water.

### Viral injections and implants

#### General procedure

Mice were anesthetized with isoflurane (1.5%) and given preemptive analgesia (extended release buprenex, 1mg/kg, and meloxicam, 5mg/kg, s.c.). After hair removal and sterilization of the skin with 70% ethanol and betadine, the mouse was placed into a stereotaxic frame (51725D, Stoelting). The temperature of the mice was maintained at 36°C with a feedback-controlled heating pad (ATC2000, World Precision Instruments). For viral injections, a micropipette attached on a Quintessential Stereotaxic Injector (QSI 53311, Stoelting) was used (see *Axonal Tracing, Axonal Imaging* and *Somatic imaging with DREADDs* for details). The pipette was held in place for 5 min after each viral injection before being slowly retracted from the brain. Postoperative analgesics (meloxicam 5mg/kg, s.c.) were given 18-24h after the surgery, and recovery was monitored for a minimum of 72 h after surgery.

#### Retrograde tracing

A small craniotomy was drilled above the right Primary Visual Cortex (AP: -3.5, ML: - 2.5, DV: -0.3) and 0.2μl of retroAAV-hSyn-GFP (50465-AAVrg, Addgene, 7×10¹² vg/mL) was injected at a rate of 0.05μl/min. Retrograde transport and virus expression was allowed for 2 weeks before the mouse was perfused for tissue collection.

#### Axonal tracing

A small craniotomy was drilled above the right Anterior Cingulate Cortex (AP: +1, ML: -0.3, DV: -0.9) and 0.2μl of AAV1-CAG-tdTomato (59462-AAV1, Addgene, 5×10¹² vg/mL) was injected at a rate of 0.05μl/min. A second craniotomy was drilled above the Orbitofrontal Cortex (AP: +2.45, ML: -1.1, DV: - 1.75) and 0.2μl of AAV5-CAG-GFP (37825-AAV5, Addgene, 7×10¹² vg/mL) was injected at a rate of 0.05μl/min. Virus expression was allowed for 4 weeks before the mouse was perfused for tissue collection.

#### Anterograde tracing

A small craniotomy was drilled above the right Anterior Cingulate Cortex (AP: +1, ML: -0.3, DV: -0.9) or Orbitofrontal Cortex (AP: +2.45, ML: -1.1, DV: -1.75) and 0.3μl of AAV8-hsyn-SytNB-ALFAtag-BACEcs-ATLAsmyc-Cre (Center for Neural Circuit Mapping, University of California, Irvine, 1×10¹^3^ gc/mL) was injected at a rate of 0.05μl/min. A second craniotomy was performed over the Visual (AP: -3.5, ML: -2.5, DV: -0.4) and the Motor Cortex (AP: +0.5, ML: -1.5, DV: -0.4) and 0.3μl of AAV.PHP.eB-CAG-DIO-tdTomato (28306-PHPeB, Addgene, 1×10¹3 vg/mL) was injected at a rate of 0.05μl/min. Virus expression was allowed for 4 weeks before the mouse was perfused for tissue collection.

#### Axonal Imaging

A small craniotomy was drilled above the Anterior Cingulate Cortex (AP: +1, ML: -0.3, DV: -0.9) or Orbitofrontal Cortex (AP: +2.45, ML: -1.1, DV: -1.75) and 0.2μl of AAV1-hSyn-axon-GCamP6s (111262-AAV1, Addgene, 7×10¹² vg/mL) was injected at a rate of 0.05μl/min. A round 3mm craniotomy was performed over the Visual (AP: -3.5, ML: -2.5) or Motor Cortex (AP: +0.5, ML: -1.5). A cranial window made of three round coverslips (CS-5R, 1×5 mm diameter; CS-3R, 2×3 mm diameter; Warner Instruments) glued together with UV-cured adhesive (catalog #NOA 61, Norland) was implanted over the craniotomy and sealed with dental cement (C&B Metabond, Parkell). For head fixation during the calcium imaging, a head plate was also affixed to the skull using dental cement (C&B Metabond, Parkell).

#### Somatic imaging with DREADDs

A small craniotomy was drilled above the Visual Cortex (AP: -3.5, ML: - 2.5, DV: -0.3) and two adjacent injections of 0.2μl retroAAV-Ef1a-Cre (55636-AAVrg, Addgene, 7×10¹² vg/mL) were injected at a rate of 0.05μl/min. After the retraction of the injection needle the craniotomy was filled with bone wax (Medline). A small craniotomy was then drilled above the Anterior Cingulate Cortex (AP: +1, ML: -0.3, DV: -0.9) or ORB (AP: +2.45, ML: -1.1, DV: -1.75) and 0.2μl AAV-hSyn-DIO-hM4D(Gi)-mCherry (44362-AAV5, Addgene, 7×10¹² vg/mL) was injected at a rate of 0.05μl/min. The skin incision on top of the skull was then closed with sutures. Two weeks after the first surgery, a second larger (3mm) craniotomy was made over the Visual Cortex, and four small injections of a mixture of 1:20 AAV-hSyn-GCamP7f (104488-AAV9, Addgene, 1×10¹³ vg/mL) and 1:5 AAV1-Ef1a-fDIO-tdTomato (128434-AAV1, Addgene, 1×10¹³ vg/mL) was made into the craniotomy. Thereafter, a cranial window made of three round coverslips (CS-5R, 1×5 mm diameter; CS-3R, 2×3 mm diameter; Warner Instruments) glued together with UV-cured adhesive (catalog #NOA 61, Norland) was implanted over the craniotomy and sealed with dental cement (C&B Metabond, Parkell). For head fixation during the calcium imaging, a head plate was also affixed to the skull using dental cement (C&B Metabond, Parkell).

### Anatomical and Histological analysis

#### General procedure

Following an overdose of isoflurane, all mice were transcardially perfused with 1x PBS followed by 4% paraformaldehyde in PBS. Brains were left over night in 4% PFA and washed three times in PBS the following day.

#### Axonal tracing

The whole brain was cut coronally at a thickness of 50μm using a vibratome (VT1200S, Leica). Every other section was thereafter collected. The collected sections were placed free-floating in 1× PBST (0.3% Triton-X in 1× PBS) for 1 h, and subsequently incubated with primary antibodies at a concentration of 1:1000 in PBST (anti-RFP: rabbit, 600-401-379, Rockland and anti-GFP: chicken, GFP-1020, Aves) overnight in room temperature. The following day the sections were washed three times in PBST and incubated with secondary antibodies (anti-Chicken Alexa Fluor 488: Donkey, 703-545-155, Jackson ImmunoResearch and anti-Rabbit Alexa Fluor 594: Donkey, 711-585-152, Jackson ImmunoResearch) at a concentration of 1:1000 in PBST for 3-5h in room temperature. The sections were consecutively washed with 1× PBST, 1× PBS and 1× PBS (10 min each). Vibratome cut sections were mounted on glass slides (Superfrost Plus, Thermo Scientific). All sections were coverslipped (Thermo Scientific) using 50:50 glycerol:1× PBS. Tiled whole-brain images were acquired of both the green and red channel using a Leica TCS SP8 confocal microscope using a 10X / 0.40 NA objective.

For the whole-brain axonal mapping of the two projections, the axonal labeling (each fluorescent pixel) was segmented out using a custom macro in ImageJ and the coordinates of each axon-labeled pixel was saved. Each section was thereafter manually assigned an AP coordinate and mapped to the Allen Brain Atlas (2011 version^20^) using a custom R-script utilizing the WholeBrainSuite R package^50^. The segmented axons were then mapped back onto the section and a pixel density value (axon-labeled pixels / total pixels) and axon density (μm^2^_axon_/μm^2^_total area_) for each brain region and layer were produced for each brain section. To avoid an overrepresentation of axonal density at each injection site, the segmentation method was modified to only extract pixels representing cell bodies without the surrounding neuropil.

#### Projection tracing and cell type labeling

The whole brain was cut coronally at a thickness of 80μm using a vibratome (VT1200S, Leica). 6-8 brain sections were selected per region including the Primary Motor Cortex and Primary Visual Cortex. The collected sections were placed free-floating in blocking buffer at room temperature overnight (2% Triton-X, 10% Normal Donkey Serum, 0.04% NaN_3_ in 1x TBS). The following day, the brain sections were transferred to primary antibody solution (1:1000 anti-PV: guinea pig, AB_572259 ImmunoStar; 1:200 anti-SST: rat, MAB354, Millipore Sigma, 1:200 anti-VIP: rabbit, AB_572270, ImmunoStar in blocking buffer) and incubated 6-8h in room temperature, followed by 2 nights in a cold room, followed by 6-8h at room temperature. The sections were washed 3 times for 3 minutes followed by 3 times for 15 minutes each in blocking buffer. The brain sections were then transferred to secondary antibody solution at a concentration of 1:1000 antibody to blocking buffer. (anti-Guinea Pig DyLight 405: Donkey, 706-475-148, Jackson ImmunoResearch; anti-Rat Alexa Fluor 647: Donkey, 712-605-153, Jackson ImmunoResearch; anti-Rabbit Alexa Fluor 488: Donkey, 711-545-152, Jackson ImmunoResearch) and incubated overnight at room temperature. The sections were again washed 3 times for 3 minutes followed by 3 times for 15 minutes each in blocking buffer. Finally, the sections were washed with 1× TBS and 1× PBS for 10 min each. All washes and incubation periods were performed on a shaker. Vibratome cut sections were mounted on glass slides (Superfrost Plus, Thermo Scientific). All sections were coverslipped (Thermo Scientific) using 50:50 glycerol:1× PBS.

Single tile images were acquired using a Leica TCS SP8 confocal microscope using a 10X / 0.40 NA objective. The XY image dimensions were 1551 by 1551µm with a Z range of 15 to 20 µm across 5 Z planes. For analysis, the images were maximum projected along the Z planes for each channel. The cell ROIs were manually labelled in the tdTomato channel using FIJI. To determine overlap of tdTomato cells with interneuron cell types, mean intensity measurements were made of the tdTomato-based ROIs across each of the channels (PV/SST/VIP). If the mean intensity of an ROI was 2 standard deviations above the pooled ROI average intensity of the respective channel (PV/SST/VIP), the cell was identified as a positive overlap. A positive overlap indicates that the identity of the tdTomato cell is PV, SST, or VIP respectively. The ROI measurements were pooled by animal and thus the threshold for interneuron (PV/SST/VIP) identification was determined per animal. The overlap/cell identity results are manually inspected for quality control.

### Behavioral setup

Mice were head-fixed on a behavioral rig, placed on top of a running wheel (Bio-Serv) attached to a rotary encoder (LPD3806-600BM-G5-24C). A monitor (15 by 9cm, 800 by 600-pixel resolution) was placed perpendicularly 14cm in front of the eye contralateral to the imaged hemisphere. Drifting gratings (0.04 cycles per degree, 2 cycles per second, 8 directions) or natural movies were displayed to the mouse at three contrasts (4%, 16%, 64%), while the mouse was free to run on the running wheel. In some imaging sessions, a mild air-puff was delivered 2 seconds before the onset of 90_°_ gratings or movie number 1, on alternating blocks of stimuli presentations. The delivery of an air puff was through a small tube 3 cm away from the whisker pad and eye ipsilateral to the imaged hemisphere (compressed air at 40 psi for 0.3 s) and was controlled through a solenoid valve (003-0137-900, Parker) and microcontroller board (Arduino UNO). Timing of stimuli and air puff as well as collection of running speed was controlled with a custom MATLAB script, with visual stimuli being generated through Psychtoolbox, and running wheel rotation and air-puff valve openings being logged through two microcontroller boards (Arduino UNO). The face and pupil of the mouse was recorded with a Thorlabs CMOS camera (CS165MU - Zelux®, Thorlabs) with an infrared filtered lens (#59-871, 25mm C Series Fixed Focal Length Lens, Edmund Optics). Infrared illumination at 780nm was provided by a light-emitting diode array light source (Thorlabs LIU780A). Video acquisition of the face was performed at 20 Hz by a custom MATLAB script. The presentation of visual stimuli, pupil imaging, and calcium imaging were aligned via 5-volt square wave pulses generated by the computer controlling stimulus timing via a National Instruments data acquisition device (National Instruments BNC-2110 Terminal Block).

### Two-photon imaging parameters

Two weeks after the mice were implanted with a cranial window, the fluorescent signal (axon-GCaMP6s or GCaMP7f) was imaged using resonant-galvo scanning with a Prairie Ultima IV two-photon microscope system. A XLUMPlanFL N 20× 1.00NA (Olympus) objective was used for single plane imaging with a field of view size of 588×588μm. Axonal activity was recoded at a 2x digital zoom and somatic activity was recorded with either 1.5x or 1x digital zoom, at a 512×512 pixel resolution. Two-photon excitation of GCaMP at a wavelength of 920 nm was provided by a tunable laser (Insight X3+, Spectra-Physics). Power at the objective ranged from 10 to 30 mW depending on depth and expression levels. Images were acquired at a frequency of 30.2Hz and every 4 frames were averaged for axonal imaging and every other frame for somatic imaging. Axonal activity recording was performed on a new field of view every day at a depth ranging from 10-150um from the brain surface. Somatic activity imaging was performed on the same set of neurons across two days (one saline day, and one CNO day).

### Preprocessing of behavioral and imaging data

#### Pupil size

Videos captured of the face of the mouse was used to post-hoc extract 8 xy-coordinates surrounding the pupil using Deep Lab Cut^51^. ∼500 frames were manually labeled (∼20 frames/ video) and a resnet v1 50-based convolutional neural network was trained to predict the location of the 8 markers for 50,000 iterations. A new network was trained for each experimental cohort. Once the network was trained, it was used to place coordinates on unlabeled frames / videos and the quality of the labeling was manually evaluated by observing at least 10 labeled videos. The distance of each xy-coordinate to the corresponding xy-coordinate across the pupil was calculated, corresponding to the diameter of the pupil in pixels. This was repeated for the 4 pairs of coordinates labeled and averaged to provide a more stable and representable pupil diameter across the entire session. Any major outliers in label position (a label suddenly jumping away from the eye) was removed post-hoc by removing values above the 99^th^ percentile of pupil values.

#### Face movements

The same videos used to extract pupil diameter information was used to extract face movement information. The videos were processed using FaceMap^52^ extracting the absolute motion energy (abs(current_frame - previous_frame)) for the entire face of the mouse.

#### Axonal imaging

2-photon axonal activity imaging was performed at a framerate of 30.2 Hz and every 4 frames was averaged resulting in a framerate of 7.55 Hz. The tiff timeseries was motion corrected twice using the Template Matching plugin in Fiji (ImageJ). The imaging sessions were motion corrected a third time in Suite2p^24^ and ROIs extracted.

The detection of axonal ROIs in Suite2p was without boundaries of size and manually curated afterwards. The fluorescence signal from the ROI was taken as F_ROI_ = F_ROIraw_ − 0.8 ∗ F_neuropil_. For computing ΔF/F (ΔF/F= (F−F_0_)/F_0_) for each ROI, F_0_ was either calculated as the mean value of F over the entire session for analysis of continuous variables, (e.g., running, face movements) or the mean value of the immediate baseline (1-2 sec) preceding the onset of an event, for analysis regarding discrete events (e.g., a visual stimulus).

In order to cluster boutons that may potentially be from the same neuron, we performed pairwise correlations between every ROI. We then performed hierarchical clustering based on these pairwise temporal correlations. The number of clusters was determined empirically for each session by iteratively testing multiple distance thresholds to satisfy a cluster correlation threshold where 1) The minimum within/intercluster correlation was >0.2 and 2) at least to times of that of the 85^th^ percentile between/inter-cluster correlation. To ensure this did not result in any false positives, 3) the same distance threshold applied to a shuffled dataset should not have any statistical difference between the between and within cluster correlations. We weighted boutons of each cluster by the signal-to-noise ratio of the ROI, taking a weighted average of fluorescence signals to represent bouton clusters (referred to as axons throughout the paper).

#### Somatic imaging

2-photon somatic activity imaging was performed at a framerate of 30.2 Hz and every other frame was averaged resulting in a framerate of 15.1 Hz. Motion correction was performed within Suite2p and ROI detection was performed without size constraints and manually curated. Somas imaged across days were identified using ROIMatchPub and manually confirmed. The fluorescence signal from each soma was taken as F_ROI_ = F_ROIraw_ − 0.8 ∗ F_neuropil_. For computing ΔF/F (ΔF/F= (F−F_0_)/F_0_) for each soma, F_0_ was either calculated as the mean value of F over the entire session for analysis of continuous variables, (e.g., running, face movements) or the mean value of the immediate baseline (1-2 sec) preceding the onset of an event, for analysis regarding discrete events (e.g., a visual stimulus).

### DREADD manipulations

In the chemogenetic DREADD experiments, mice were injected with 1mg/kg Clozapine-N-oxide intraperitoneal (i.p.) or with saline (0.9% NaCl) of the same volume of the drug administration. Mice were imaged at least 30min after the i.p. injection. Chemogenetic constructs were only expressed unilaterally in these experiments.

### Analysis

#### Analysis of visual responses in axons or somas

To determine if an axon or soma was visually responsive, we compared the mean value of activity 1 second pre-stimulus with the mean activity during stimulus presentation (1 second for gratings, 2 seconds for movies). A paired t-test was then performed between pre and post responses (at 3 contrasts, 10 trials/values for each contrast) for each direction of grating or individual movie. Axons or somas with a p-value < 0.00625 for one or several directions of drifting grating or with a p-value < 0.01 for one or several movies were considered visually responsive. If an axon or soma was significantly responsive to multiple directions or movies, the direction or movie with the highest response amplitude was considered the preferred stimulus.

To compare single-cell visual responses in the presence or absence of PFC modulation, only neurons that were tracked across days were used (matched neurons). From the population of matched neurons, only neurons that kept their visual responsiveness and at least one preferred direction/movie across imaging day were kept for further comparative analysis.

Orientation tuning curves of matched neurons were created by identifying the preferred stimulus of a neuron on the saline day (highest amplitude of response), and aligning the activity from the cno day to the same preferred direction. Response amplitude for each direction was the mean ΔF/F activity during visual stimulus presentation, averaged across all presentations of that direction. The direction selectivity index was also computed as:

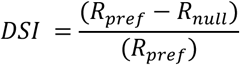

Where the R_pref_ and R_null_ are responses at the preferred (pref) and preferred + 180° (null) orientation respectively.

#### Behavioral state

For a visual response to be classified as occurring during a discrete behavioral state the mouse needed to be moving/aroused above the set threshold (defined for each panel in the figure ledged) for the entire trial, 1 sec before, during visual stimuli, and 1 second after visual stimuli.

#### Decoding stimulus identity and contrast

We built linear support vector machine (SVM) classifiers using the *fitceco* function in MATLAB. All population decoding was performed for each session. For the training and testing dataset, the number of trials in each condition (eight directions and three contrasts) was matched to prevent bias for training classifiers. We used three-fold cross-validation by leaving a 33% subset of trials for prediction to avoid overfitting. This procedure was repeated 50 times. To avoid overestimating the weights, we applied Lasso regularization using a *templateLinear* function in MATLAB. Hyperparameter such as *+* regularization weight was determined by optimization to minimize loss of validation dataset in a grid search manner (searched range 10^−6^ – 10^−1^).

We used the mean ΔF/F response during stimulus presentation (1 second for gratings, 2 seconds for movies) for each individual neuron, respectively. Classifier performance on each iteration was estimated by averaging decoding accuracies across the three folds. Final decoding accuracy was determined by averaging these mean accuracies across all iterations. To obtain cross-validated single-trial accuracy values, we averaged the binary prediction accuracy of each trial across all cross-validation folds and iterations in which it was held out as part of the test set. To determine if the decoder was informative above chance, we shuffled labels for the test data and trained and tested decoder to assess the decoding accuracy.

For statistically assessment of the decoding accuracy between sessions, we trained and tested decoders using subset of population of neurons (from 10-200 neurons, at 10 neuron increments) by randomly choosing neurons in each iteration. Difference in decoding accuracy between two conditions (saline and CNO) were evaluated in multiple ways. First, by performing a 2-way ANCOVA to examine the effects of subset (number of neurons) and experimental condition (sal/cno) on the accuracy of decoding. The model included interaction terms to explore if the effect of subset size on accuracy varies by group and data/shuffled data. p-value noten in the figures are from the variability in accuracy explained by group (sal vs. cno). Second, by performing a paired t-test on the average accuracy per subset across paired recording sessions (imaging field of views). Lastly, by computing the difference in decoding accuracy for each group and subset between the data and the shuffled data, and thereafter performing a paired t-test of the differences in delta decoding accuracy between paired recording sessions and subset size.

#### Generalized Linear Model (GLM) of soma and axonal activity

To identify the visual information and behavioral variables of soma and axon, we used a GLM with linear (identity) link function. In this model, the soma and axonal activity is described as a linear sum of visual and behavioral predictors aligned to each event. The predicted soma or axonal activity *r_n_*(*t*) for a soma or axon. is described as

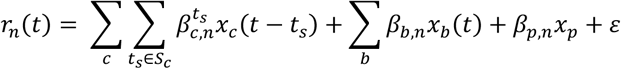

where 4 represents the direction and contrast of visual stimulus (eight direction x three contrasts), 5 represents the behavioral variables (pupil diameters, face velocity, running speed, binarized pupil diameters, and binarized running speed), *p* represents the air puff, *s_c_* represents the set of times to cover each predictor window. *β_c,n_, β_b,n,_, β_p,n_* represents the weights of visual stimulus, behavioral variables, and air puff for soma or axon .. The visual stimulus predictors cover the window -1-3 s from stimulus onset. *x_c_, x_b_, x_p_* represents the visual stimulus, behavioral variable, and air puff predictors. Each predictor coded as “1” or “0” except for behavioral variables. Behavioral variable predictors are the continuous behavioral event variables such as pupil diameter. 3 is the model bias (intercept). The values for the behavioral variables were z-scored. The soma or axonal activity was binned into 0.2seconds.

To estimate the optimal weights for each soma or axonal activity without overfitting, the *lassoglm* function in MATLAB with tenfold cross-validation of the training set was used with a lasso regularization according to the value of a selected parameter +, which represents regularization coefficients. The value of + in the lassoglm function was set to be 10^−3^. Model performance was assessed for the test dataset by quantifying explained variance (R^2^).

To determine the contribution of each variable to soma or axonal activity, we fitted the model using full predictors (full model) and predictors in which the target predictor is set to zero within whole-time points (partial model) and calculated the explained variance 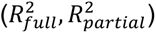 of the full and partial model. We defined relative contribution of each variable to soma or axonal activity by determining how much the performance of the partial model declined compared to full model.

In this study, proportional contribution of variable : is calculated as proportion of all variable’s contribution to the model:

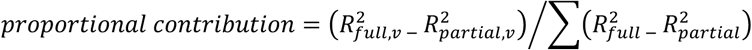

where 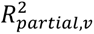 represents explained variance of partial model of *v* th variable. Negative relative contributions were set to zero (this means partial model performed better than full model).

The proportion of axons or somas with a significant contribution of the variance explained by variable : was calculated by performing a t-test between the 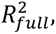, and 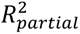 across the 10-fold cross validation, correcting for multiple comparisons with Bonferroni-Holm correction.

### Statistical Methods

Statistical methods are noted in the figure legends and result section. Statistical analysis was carried out in Python 3.8.8 using the sklearn and scipy libraries.

## Notes

### Competing Interest Statement

The authors have declared no competing interest.

### Summary of Updates

Revisions to both text and figures. Additional analysis has been added to Figure 2 and 4.

