## Supplementary Figures for "Prefrontal Cortex subregions provide distinct visual and behavioral feedback modulation to the Primary Visual Cortex"

### EXTENDED DATA FIGURES

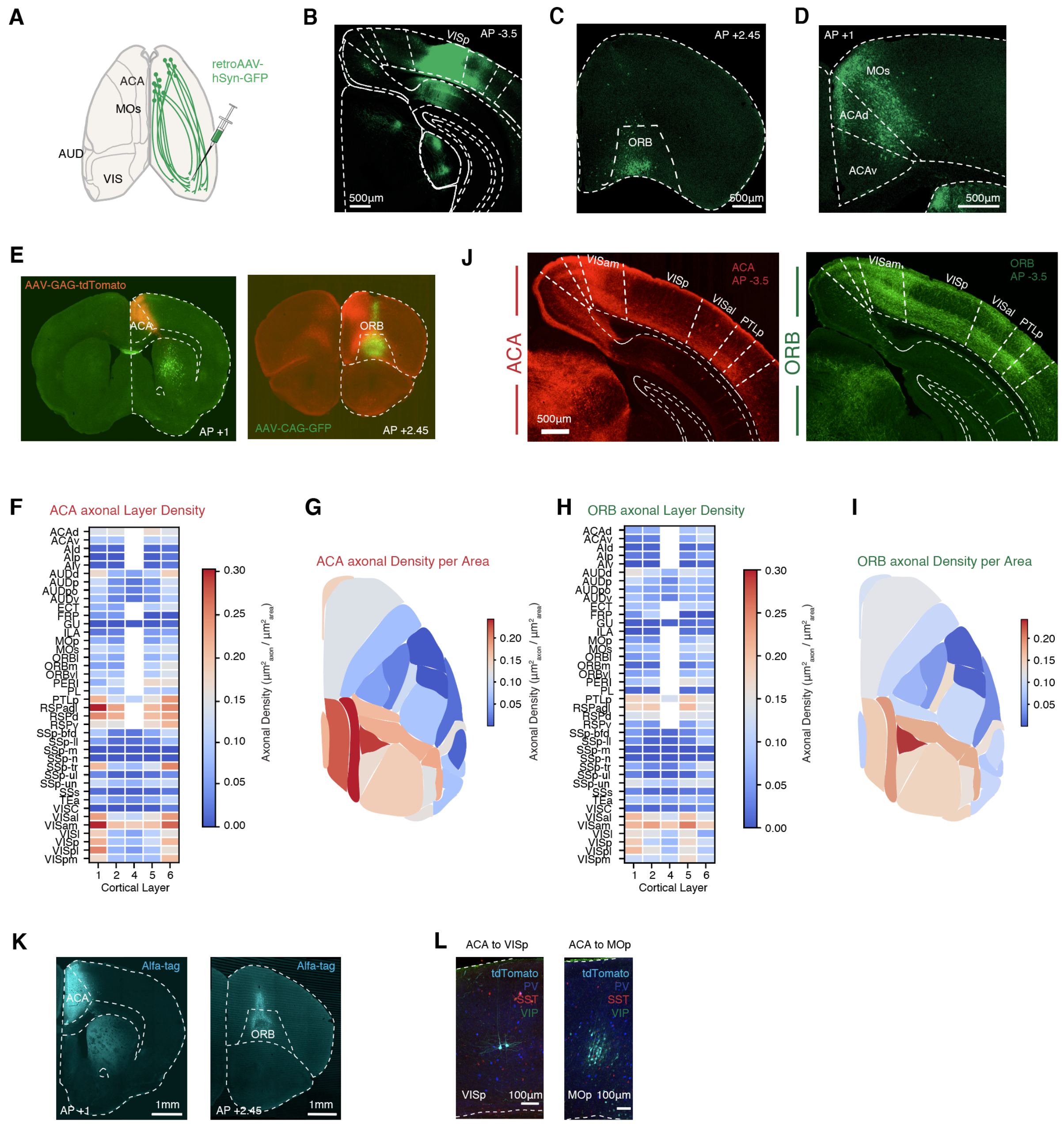


**Supplementary Figure 1. Axonal projections from the ACA and the ORB.**
**(A)** Experimental strategy. retroAAV-hSyn-GFP was injected into the VISp.
**(B)** Representative image of the injection site. The experiment was replicated in n=2 mice.
**(C)** Representative image of retrograde labeling of neurons in the ORB.
**(D)** Representative image of retrograde labeling of neurons in the ACA and the secondary Motor cortex.
**(E)** Experimental strategy of axonal labeling of ACA and ORB projection neurons (left). AAV-GAG-tdTomato was injected into the ACA (middle) and AAV-CAG-GFP was injected into the ORB (right).
**(F)** Average axonal density of ACA axons per cortical region and layer (n=4 mice, 67±20 brain sections per animal).
**(G)** Average axonal density of ACA axons per cortical region as schematic seen from above.
**(H)** Average axonal density of ORB axons per cortical region and layer (n=4 mice, 67±20 brain sections per animal).
**(I)** Average axonal density of ORB axons per cortical region as schematic seen from above.

**(J)** Representative images of ACA (red) and ORB (green) axons in the primary Visual cortex and surrounding visual areas. Scale bar 500$\mu$m. Related to images in Figure 1.
**(K)** Representative images of injection sites of AAV-ATLAS$_{Cre}$ in the ACA (left) and ORB (right). Immunohistological staining against the Alfa-tag (cyan) expressed by the virus allowed for confirmation of injection site location. Scale bar: 1mm. (Replicated in ACA: n=4 mice, ORB: n=3 mice).
**(L)** Representative images of tdTomato positive cells (cyan) in the VISp (left) and MOp (right) labeled by AAV-ATLAS$_{Cre}$ and AAV-DIO-tdTomato co-expression. Immunohistological staining of SST (red), VIP (green) and PV (blue) allowed for the quantification of interneurons targeted by ACA or ORB axons. Replicated in ACA: n= 4 mice, MOp n=32 sections, VISp n=31 sections. ORB: n= 3 mice, MOp:30 sections, VISp: 15 sections.

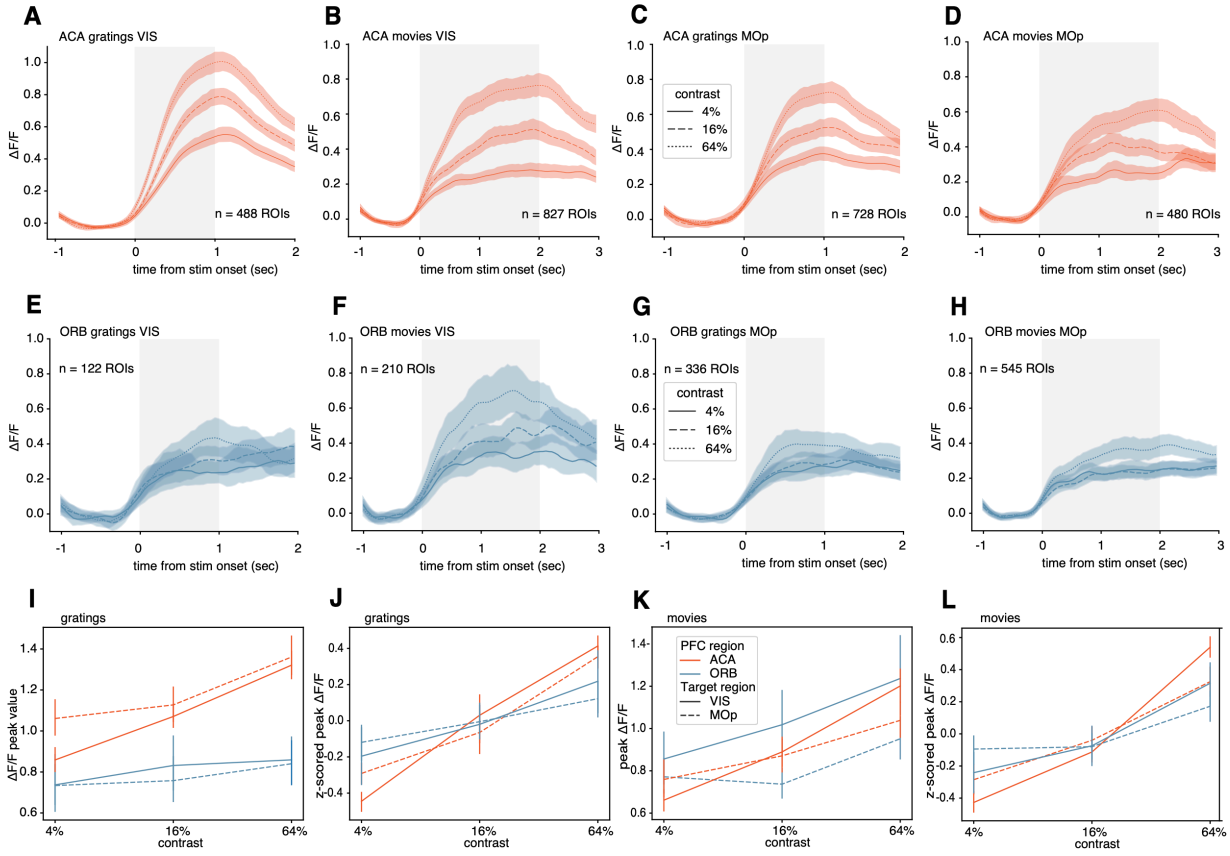


**Supplementary Figure 2. ACA, but not ORB, axonal activity scales response amplitude to contrast of visual stimuli.**

**(A-D)** Averaged population responses of all visually responsive ACA axons with an increase in ΔF/F in response to stimuli, plotted per contrast of stimuli, for each target area and stimulus type. ACA-VIS 4 mice, ACA-MOp 4 mice. Solid or dashed line: mean value, shaded area: 95% confidence interval.

**(E-H)** Averaged population responses of all visually responsive ORB axons with an increase in ΔF/F in response to stimuli, plotted per contrast of stimuli, for each target area and stimulus type. ORB-VIS 4 mice, ORB-MOp 4 mice. Solid or dashed line: mean value, shaded area: 95% confidence interval.

**(I)** Mean ΔF/F peak responses during stimuli on-time plotted per contrast. Visually responsive ACA and ORB axons in VIS and MOp, with a significant increase in ΔF/F in response to grating stimuli (paired t-test of mean ΔF/F one second before stim onset vs during stim on time, corrected for multiple comparisons with Bonferroni correction, $p < 0.00625$ for gratings) are included.

**(J)** Mean standardized ΔF/F peak responses during stimuli on-time plotted per contrast. Visually responsive ACA and ORB axons in VIS and MOp, with a significant increase in ΔF/F in response to grating stimuli (paired t-test of mean ΔF/F one second before stim onset vs during stim on time, corrected for multiple comparisons with Bonferroni correction, $p < 0.00625$ for gratings) are included. ΔF/F mean responses are z-scored across each individual axon for standardization.

**(K)** Mean ΔF/F peak responses during stimuli on-time plotted per contrast. Visually responsive ACA and ORB axons in VIS and MOp, with a significant increase in ΔF/F in response to movies stimuli (paired t-test of mean ΔF/F one second before stim onset vs during stim on time, corrected for multiple comparisons with Bonferroni correction, $p < 0.01$ for movies) are included.

**(L)** Mean standardized ΔF/F peak responses during stimuli on-time plotted per contrast. Visually responsive ACA and ORB axons in VIS and MOp, with a significant increase in ΔF/F in response to movies stimuli (paired t-test of mean ΔF/F one second before stim onset vs during stim on time, corrected for multiple comparisons with Bonferroni correction, $p < 0.01$ for movies) are included. ΔF/F mean responses are z-scored across each individual axon for standardization.

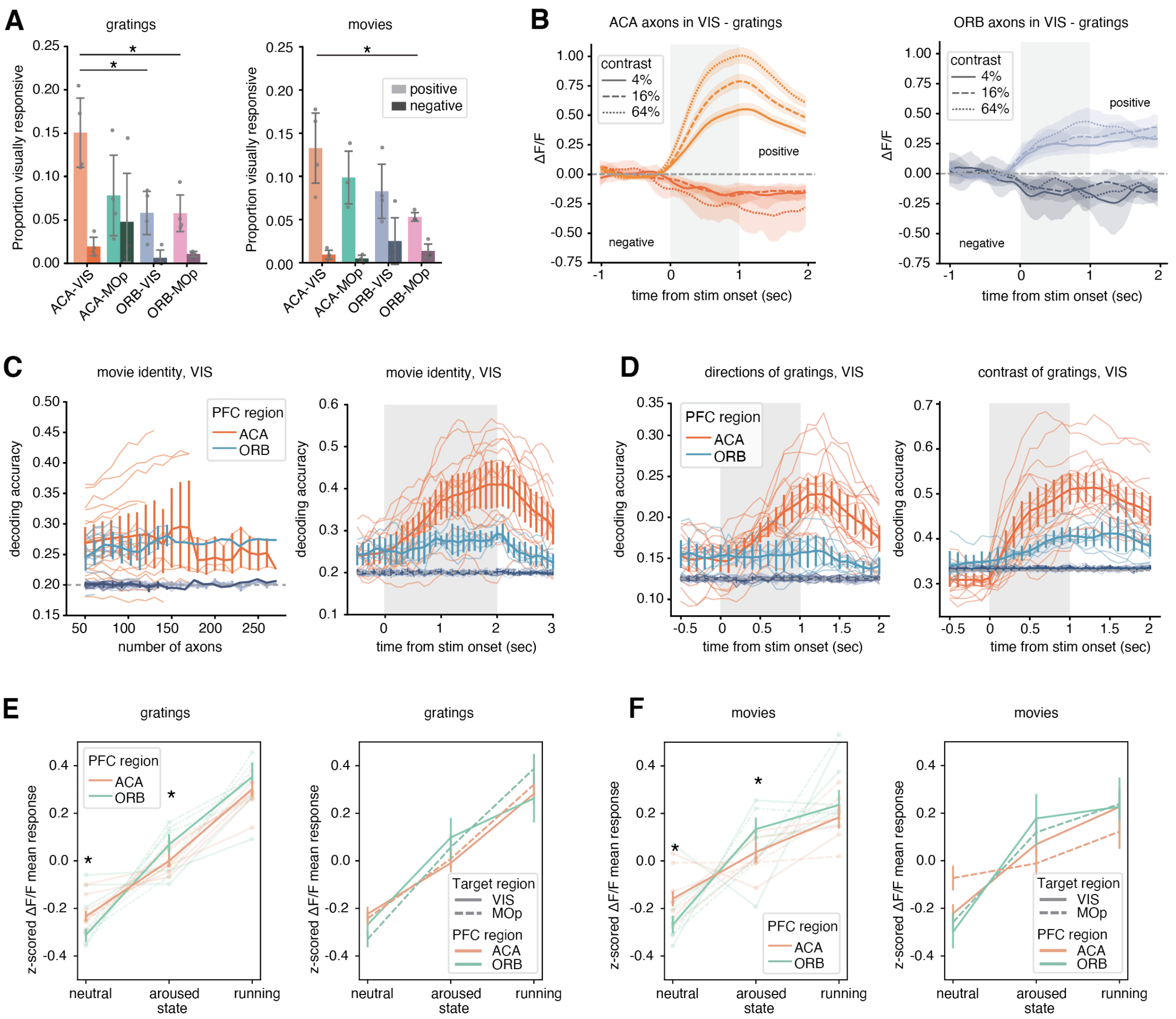


**Supplementary Figure 3. ACA and ORB axonal response amplitude to visual stimuli is modulated by behavioral state.**

**(A)** Proportion of visually responsive axons to gratings (left) and movies (right) from the ACA or the ORB in the VIS or the MOp, plotted separated by response type. Bars display mean of proportions, error bars: SEM, dots represent individual animals. Gratings, positively modulated axons: ACA-VIS vs ORB-VIS: p-value= 0.0296, ACA-VIS vs ORB-MOp: p-value= 0.029. Movies, positively modulated axons: ACA-VIS vs ORB-MOp: p-value= 0.0348. one-way ANOVA with post-hoc Tukey test for multiple comparisons of means. N= 4 mice for each group.

**(B)** Averaged population responses of all visually responsive axons with an increase or decrease in ΔF/F in response to gratings, plotted per contrast (left; ACA-VIS axons, right; ORB-VIS axons). Solid line: mean value, shaded area: 95% confidence interval.

**(C)** Left, decoding accuracy of SVM decoder of movies trained and tested on ACA (orange) or ORB (gray) axonal responses (mean ΔF/F during stimuli on-time) in the VIS, per imaging session and field of view). Right, decoding accuracy of SVM decoder of movies over time. 100 axons included per session. Decoding accuracy of shuffled ACA axonal responses (light gray) or shuffled ORB axonal responses (dark gray) plotted as comparison. Thick solid line: mean accuracy across sessions, error bars: 95% confidence interval (omitted when only one session contributed to a bin), thin lines: individual imaging sessions, dashed line: chance level.

**(D)** Decoding accuracy of SVM decoder of direction (left) and contrast (right) of gratings, over time. The decoder is trained and tested on ACA (orange) or ORB (gray) axonal responses per time bin, in the VIS, per imaging session and field of view. Decoding accuracy of shuffled ACA axonal responses (light gray) or shuffled ORB axonal responses (dark gray) plotted as comparison. Thick solid line: mean accuracy across sessions, error bars: 95% confidence interval (omitted when only one session contributed to a bin), thin lines: individual imaging sessions, dashed line: chance level.

**(E)** Mean standardized ΔF/F peak responses during stimuli on-time plotted per behavioral state. Left: Visually responsive ACA and ORB axons in VIS and MOp. Right: Visually responsive ACA and ORB axons in VIS (solid lines) or MOp (dashed lines). Axons with a significant increase in ΔF/F in response to gratings stimuli (paired t-test of mean ΔF/F one second before stim onset vs during stim on time, corrected for multiple comparisons with Bonferroni correction, $p < 0.00652$ for gratings) are included. ΔF/F mean responses are z-scored across each individual axon for standardization. Thick lines: mean standardized ΔF/F mean responses across axons, error bars: 95% confidence interval, thin lines and dots: individual animals. * < 0.01, one-way ANOVA with post-hoc Tukey test for multiple comparisons of means.
**(F)** Mean standardized ΔF/F peak responses during stimuli on-time plotted per behavioral state. Left: Visually responsive ACA and ORB axons in VIS and MOp. Right: Visually responsive ACA and ORB axons in VIS (solid lines) or MOp (dashed lines). Axons with a significant increase in ΔF/F in response to movie stimuli (paired t-test of mean ΔF/F one second before stim onset vs during stim on time, corrected for multiple comparisons with Bonferroni correction, $p < 0.01$ for movies) are included. ΔF/F mean responses are z-scored across each individual axon for standardization. Thick lines: mean standardized ΔF/F mean responses across axons, error bars: 95% confidence interval, thin lines and dots: individual animals. * < 0.01, one-way ANOVA with post-hoc Tukey test for multiple comparisons of means.

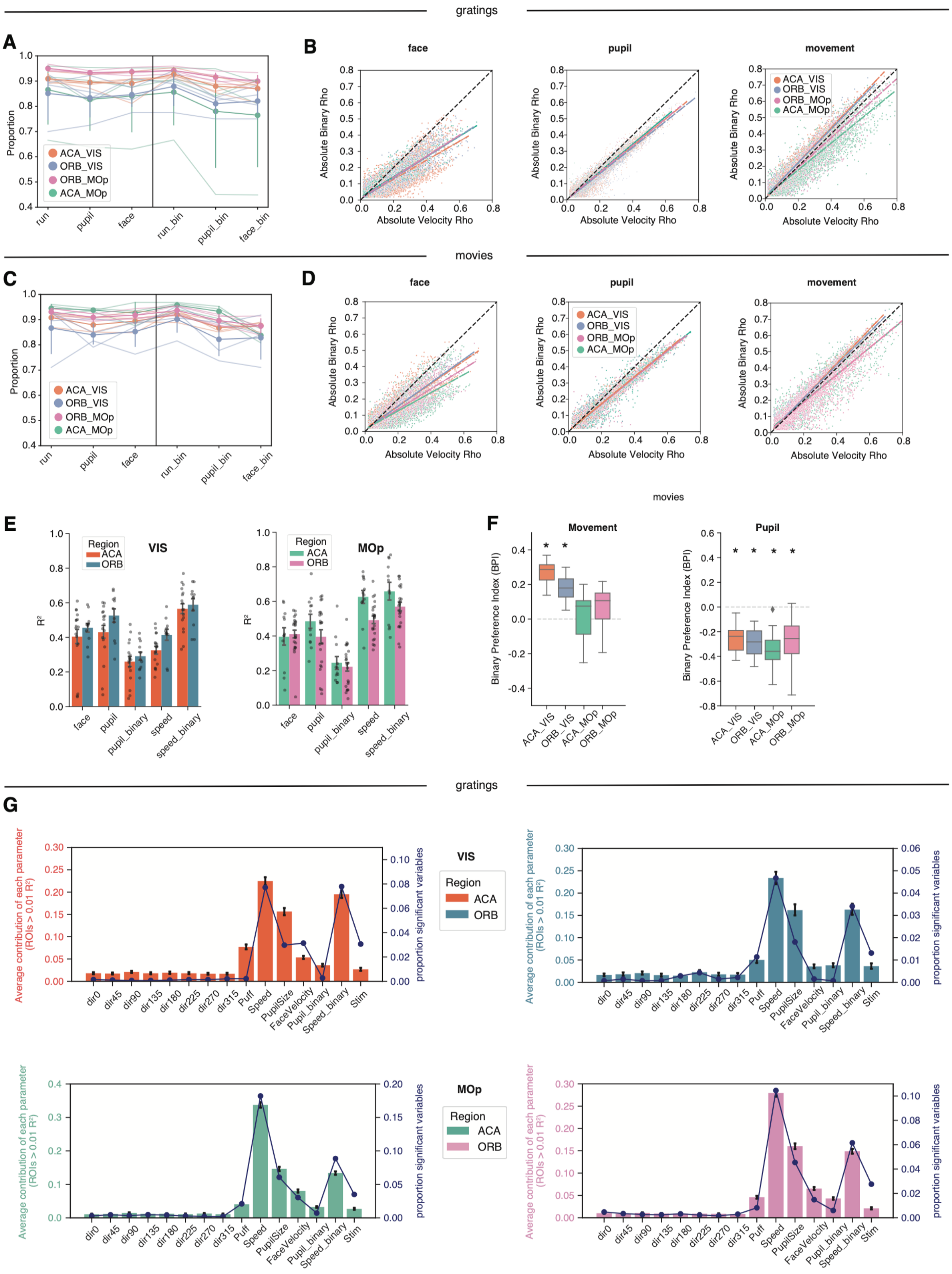


**Supplementary Figure 4. ACA and ORB axonal activity in VISp and MOp is modulated by behavior.**
**(A)** Proportion of axons recorded during grating sessions significantly (p<0.01) correlating (Spearman's correlation) to behavioral variables; running speed (cm/s), pupil size (z-score), absolute face movement (z-score), binary movement

(>0.5 cm/s), binary pupil (>1 z-score), binary face movement (>1 z-score). Circle indicates mean across all animals, color coded by source and target area. Error bars: 95% confidence interval, lines: individual animals.

**(B)** Significant ($p<0.01$) Spearman's correlation values (Rho) of each axon $\Delta F/F$ activity with the z-scored face movement against binary dilated state (left), z-scored pupil size against binary dilated state (middle), or running speed (cm/s) against binary movement state (0 or 1) (right). Each dot represents one axon color coded by the source and target area recorded during grating sessions, only axons with significant correlation to both actual and binary value are included in the plot. Dashed lines are linear regression lines for each subset of axons relationship between the actual and binary movement or pupil size.

**(C)** Proportion of axons recorded during movie sessions significantly correlating to behavioral variables; running speed (cm/s), pupil size (z-score), absolute face movement (z-score), binary movement (>0.5 cm/s), binary pupil (>1 z-score), binary face movement (>1 z-score). Circle indicates mean across all animals, color coded by source and target area. Error bars: 95% confidence interval, lines: individual animals.

**(D)** Significant ($p<0.01$) Spearman's correlation values (Rho) of each axon $\Delta F/F$ activity with the z-scored face movement against binary dilated state (left), z-scored pupil size against binary dilated state (middle), or running speed (cm/s) against binary movement state (0 or 1) (right). Each dot represents one axon color coded by the source and target area recorded during movie sessions, only axons with significant correlation to both actual and binary value are included in the plot. Dashed lines are linear regression lines for each subset of axons relationship between the actual and binary movement or pupil size.

**(E)** Proportion of variance explained (R2) of linear model of behavioral parameters (face (SVM z-scored), pupil (diameter z-scored), pupil binary value (0 or 1 if z-score > top 20$^{th}$ percentile for that imaging session), speed (cm/s), speed binary value (0 or 1 if cm/s > 0.5) predicted by the axonal activity of ACA or ORB axons in the VIS (left) or MOp (right) during movie sessions. Bars: mean R2 of all imaging sessions, error bars: SEM (68% confidence interval), dots: individual sessions. Imaging sessions included are n=17 ACA-VIS, n=11 ORB-VIS, n=12 ACA-MOp, n=22 ORB-MOp.

**(F)** Strength of correlation to actual vs binary value of movement (left) or pupil diameter (right) calculated as a binary preference index (BPI) by the variance explained for the binary versus actual value. BPI is plotted for each subset of axons based on source and target area. Each boxplot represents the quartiles of values, while the whiskers extend to show the rest of the BPI values, per imaging session, outliers represented as diamonds. * = p-value <0.05, t-test of significant difference from 0. Movement: ACA_VIS, p-value: 5.2e-12, ORB_VIS, p-value: 1.35e-05, ACA_MOp, p-value: 0.65, ORB_MOp, p-value: 0.006. Pupil: ACA_VIS, p-value: 3.09e-08, ORB_VIS, p-value: 1.10e-05, ACA_MOp, p-value: 2.54e-05, ORB_MOp, p-value: 7.97e-07.

**(G)** Proportion of parameters with significantly contributing variance explained (blue line) plotted with the average contribution of variance explained (R2) for each parameter added to a linear model predicting the activity of single axons that has at least 1% activity explained by the model. Bars: mean proportion, error bars: 95% confidence interval. Each plot is color coded by the target and source area for the axons included in the analysis.

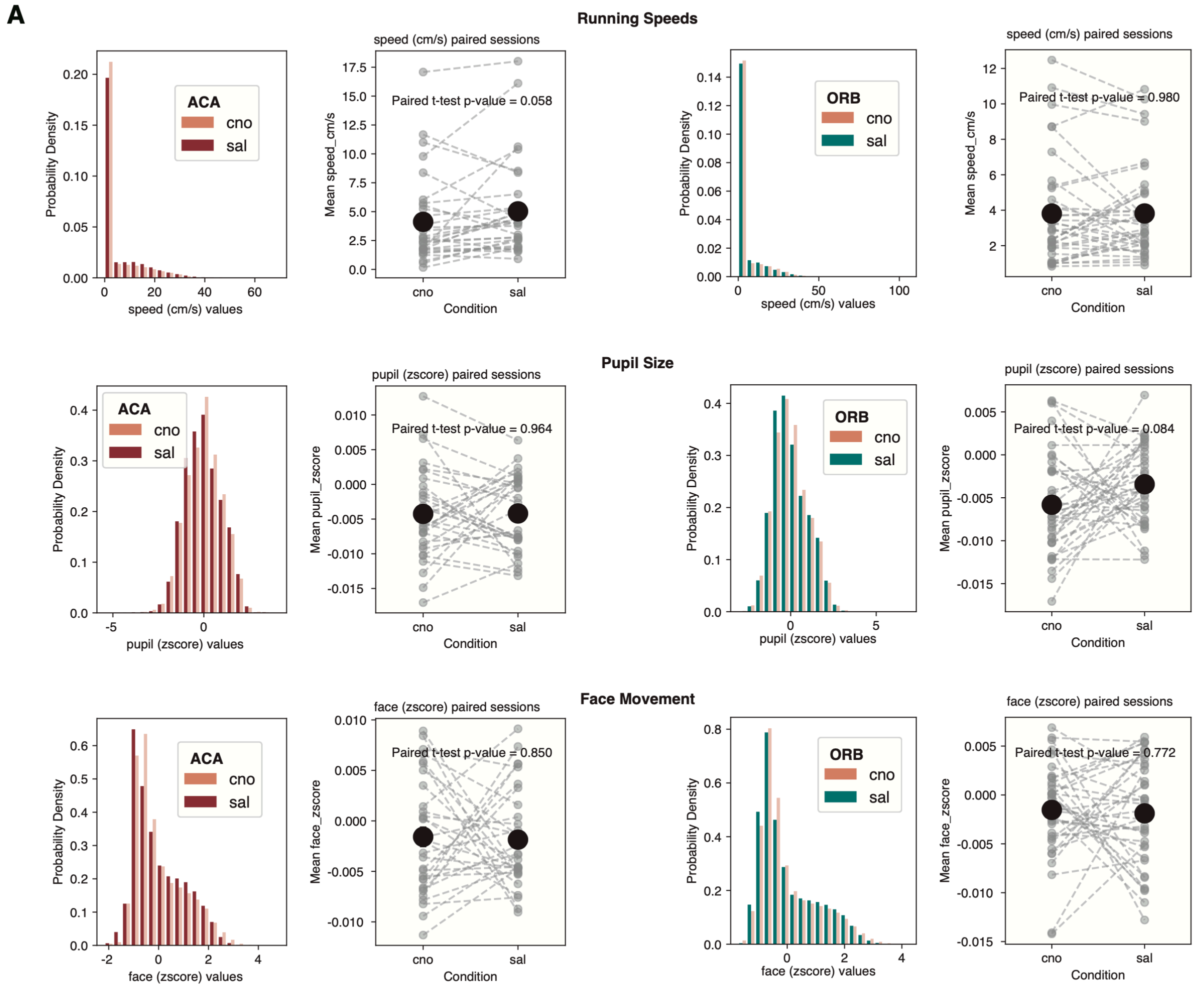


**Supplementary Figure 5. CNO or Saline injections do not change the behavior of mice during imaging sessions. (A)** Right. Probability density of speed (cm/s), pupil size (z-scored) and face movement (z-scored) values across all Saline (sal, ACA:red, ORB:green) and CNO (cno: beige) recording sessions. Left. Mean speed (cm/s), pupil size (z-scored) and face movement (z-scored) for all sessions (black) and each paired recording sessions (sal/cno) plotted as individual lines (gray). * = p-value < 0.05, paired Student's t-test. ORB: 37 paired imaging session, across n= 6 mice, ACA: 31 paired imaging session, across n= 5 mice.

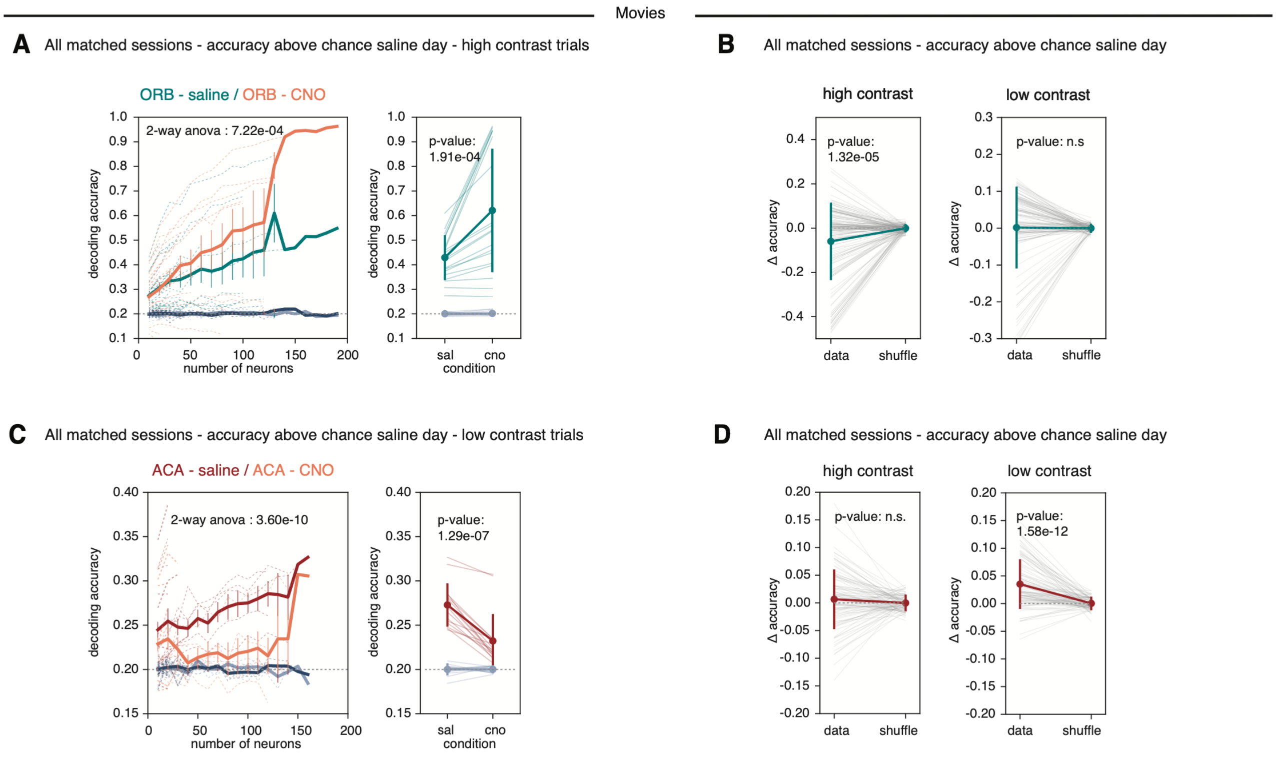


**Supplementary Figure 6. ACA modulation increases decoding accuracy of weak stimuli while ORB modulation decreases decoding accuracy of high contrast stimuli.**

**(A)** Decoding accuracy of SVM decoder of natural movies trained and tested on saline day (green-ORB) or CNO day (orange) VIS soma responses (mean ΔF/F during stimuli on-time), for high contrast trials. Decoding accuracy of shuffled soma responses saline day (light gray) or shuffled soma responses CNO day (dark gray) plotted for comparison. All paired recording sessions with a minimum accuracy above chance level on saline day included in the analysis. Thick solid line: mean accuracy across sessions, error bars: 95% confidence interval (omitted when only one session contributed to a bin), thin lines: individual imaging sessions, dashed line: chance level. ORB: 26 paired recording sessions n=6 mice. 2-way ANCOVA to examine the effects of subset (number of neurons) and experimental condition (saline/ CNO) on the accuracy of decoding. The model included interaction terms to explore if the effect of subset on accuracy varies by group and data/shuffled data. p-value from the variability in accuracy explained by group (saline vs. CNO).

**(B)** Decoding accuracy of SVM decoder of drifting gratings directions trained and tested on VIS soma responses (mean ΔF/F during stimuli on-time) on either high contrast (64%) or low contrast (16%, 4%) stimuli. Change in decoding accuracy per paired recording session (field of view) and subset (number of neurons), comparing real and shuffled data. P-value from paired t-test between real and shuffled data. High contrast trials; ORB: 29 paired recording sessions n=6 mice. Low contrast trials; ORB: 25 paired recording sessions n=6 mice.

**(C)** Decoding accuracy of SVM decoder of natural movies trained and tested on saline day (green-ORB) or CNO day (orange) VIS soma responses (mean ΔF/F during stimuli on-time), for low contrast trials. Decoding accuracy of shuffled soma responses saline day (light gray) or shuffled soma responses CNO day (dark gray) plotted for comparison. All paired recording sessions with a minimum accuracy above chance level on saline day included in the analysis. Thick solid line: mean accuracy across sessions, error bars: 95% confidence interval (omitted when only one session contributed to a bin), thin lines: individual imaging sessions, dashed line: chance level. ACA: 22 paired recording sessions n=5 mice. 2-way ANCOVA to examine the effects of subset (number of neurons) and experimental condition (saline/ CNO) on the accuracy of decoding. The model included interaction terms to explore if the effect of subset on accuracy varies by group and data/shuffled data. p-value from the variability in accuracy explained by group (saline vs. CNO).

**(D)** Decoding accuracy of SVM decoder of drifting gratings directions trained and tested on VIS soma responses (mean ΔF/F during stimuli on-time) on either high contrast (64%) or low contrast (16%, 4%) stimuli. Change in decoding accuracy per paired recording session (field of view) and subset (number of neurons), comparing real and shuffled data. P-value from paired t-test between real and shuffled data. High contrast trials; ACA: 25 paired recording sessions n=5 mice. Low contrast trials; ACA: 22 paired recording sessions n=5 mice.

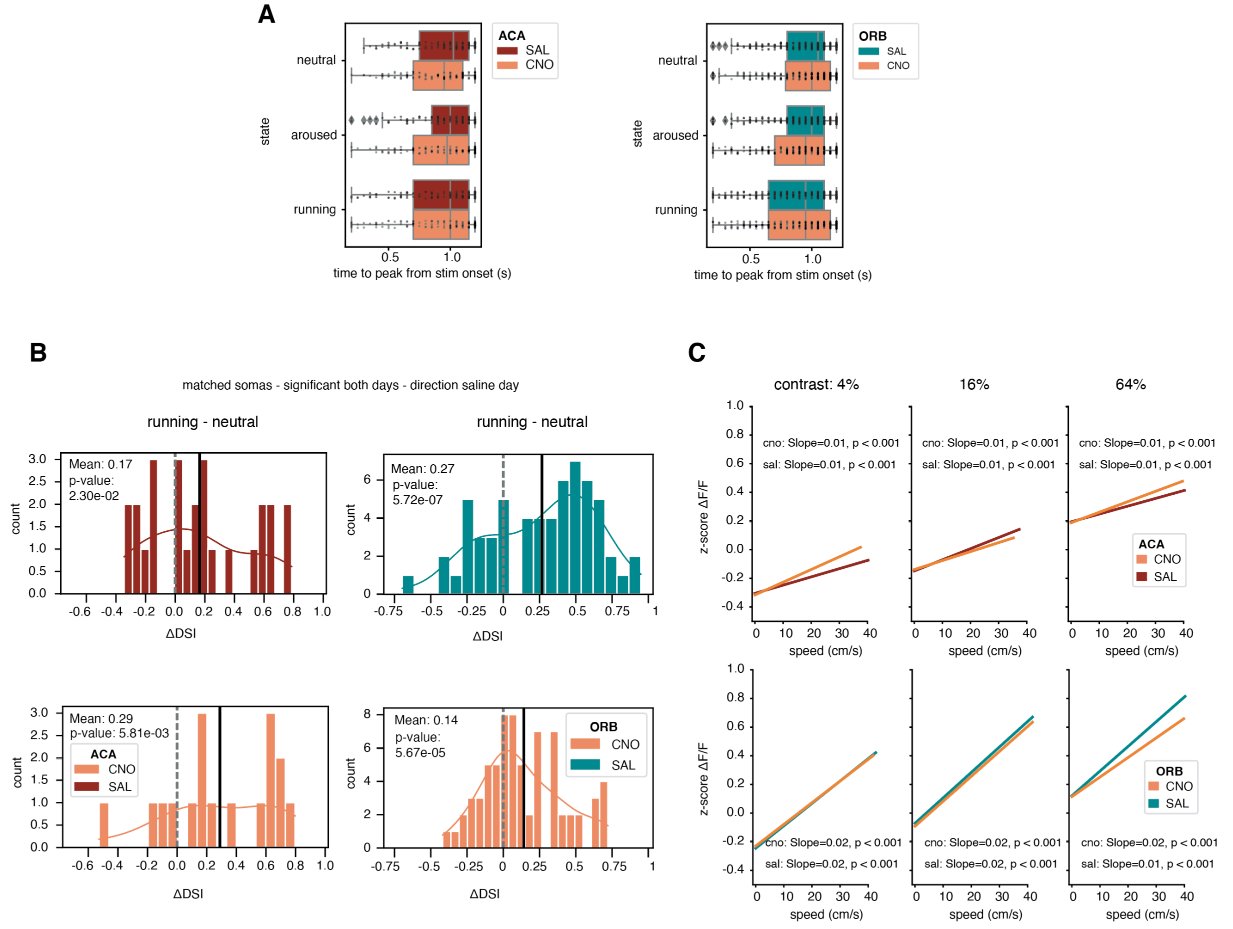


**Supplementary Figure 7. ACA and ORB modulation of visually responsive neurons relate to behavioral state.** **(A)** Boxplot of peak amplitude of visual responses of matched VISp neurons to their preferred direction(s) across behavioral states, with or without CNO inhibition of ACA (left) or ORB (right). For each neuron, the time to peak response was calculated as the time point (in seconds) at which the average ΔF/F trace reached its maximum within the stimulus window, computed separately for each behavioral state (neutral, aroused, running) and condition (saline, CNO). The horizontal box plots show the distribution of time-to-peak values across neurons for each state and condition. Individual neurons are overlaid as points. Box plots indicate the median (line), interquartile range (box), and whiskers extending to 1.5× the IQR. Statistical comparisons between saline and CNO time-to-peak values were performed separately for each state using the Wilcoxon signed-rank test (paired, two-sided), with Bonferroni correction applied for multiple comparisons across the three states. All comparisons are non-significant. ACA-DREADDs 132 neurons from 5 mice; ORB-DREADDs 328 neurons from 6 mice.

**(B)** Histogram of values of change of Direction Selectivity Index (ΔDSI) from a neutral state to a running state (top 20 percentile pupil events). ΔDSI values stem from matched neurons that were significantly visually responsive on both days to the same direction(s), aligned to the preferred direction on the saline day. Dashed line indicates 0, and the solid line is the mean of ΔDSI values plotted. p-value from one-sampled t-test against 0. ACA-DREADDs saline: 26 neurons CNO: 11 neurons from 5 mice. ORB-DREADDs saline: 61 neurons CNO: 46 neurons from 6 mice.

**(C)** Linear regression lines fit to z-scored ΔF/F single trial activity against running speed (cm/s), per condition (saline/CNO) and contrast. Single trial activity is included from matched neurons that were significantly visually responsive on both days to the same direction(s). ΔF/F mean responses for stimuli on-time per trial are z-scored across each neuron and condition. The slopes and significance levels are determined using Ordinary Least Squares (OLS) regression. Top: ACA-DREADDs 152 neurons from 5 mice, total of 49290 trials, 24737 CNO trials and 24553 saline trials. Bottom: ORB-DREADDs 380 neurons, total of 120652 trials, 60336 CNO trials and 60316 saline trials.

**A**

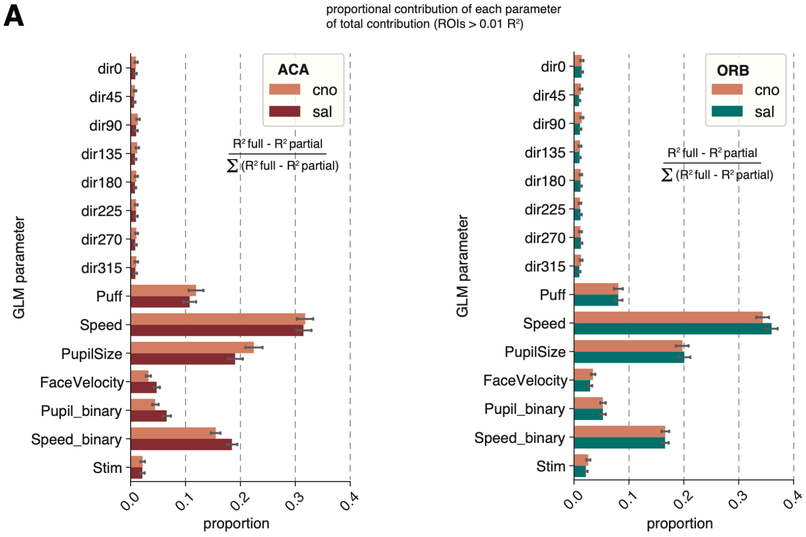


**Supplementary Figure 8. ACA and ORB modulation does not alter the proportional contributions of behavioral parameters of a generalized model of VIS soma activity.**
**(A)** Variance explained ($R^2$) of each parameter added to a linear model predicting the activity of single axons that has at least 1% activity explained by the model. Contribution of each parameter plotted as the proportional contribution of the total contribution explained per soma, plotted for each experimental condition (saline: ACA-red, ORB-green / CNO: orange). Bars: mean proportion, error bars: 95% confidence interval. ACA-saline 1459 somas, 26 imaging sessions, 5 mice. ORB-saline 2637 somas, 38 imaging sessions, 6 mice. ACA- CNO 1462 somas, 26 imaging sessions, 5 mice. ORB- CNO 2477 somas, 38 imaging sessions, 6 mice.

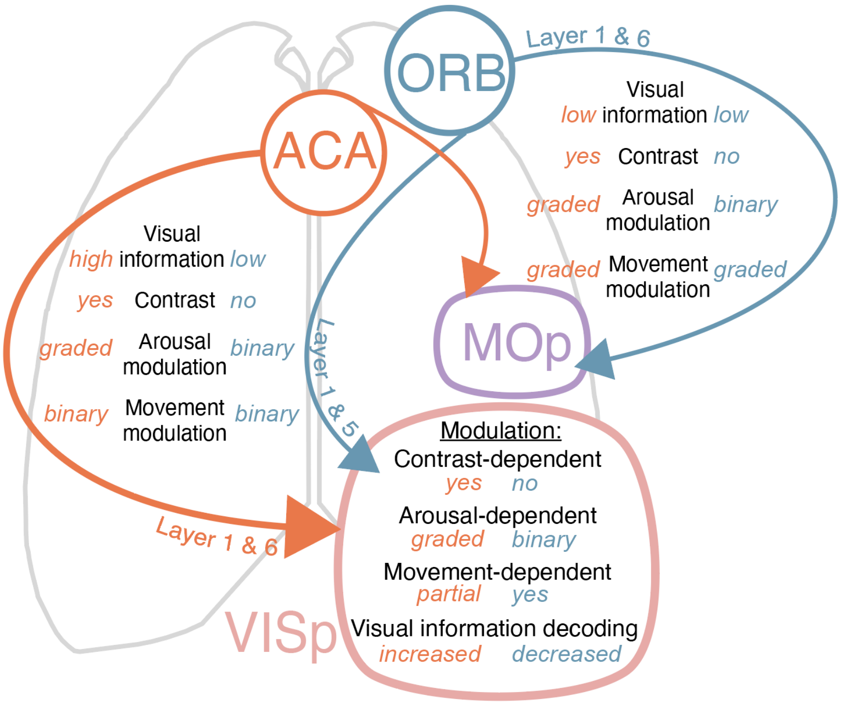


**Supplementary Figure 9. Schematic representation of ACA and ORB feedback modulation onto VISp and MOp.** ACA provides contrast-dependent visual information modulated by graded levels of arousal to VISp. Both ACA and ORB provide stronger representation of visual information to VISp compared to MOp. ACA feedback enhances visual encoding in VISp during arousal states. ORB feedback links high arousal and movement modulation to VISp activity at the expense of visual encoding. Feedback from ACA and ORB is thus specialized at both the source regions and their targets, enabling each region to selectively shape the activity of VISp neurons.
